# Assembly properties of *Spiroplasma* MreB involved in swimming motility

**DOI:** 10.1101/2023.01.26.525654

**Authors:** Daichi Takahashi, Makoto Miyata, Ikuko Fujiwara

**Author notes:** To whom correspondence should be addressed: Ikuko Fujiwara. **Competing Interest Statement**: The authors declare that they have no conflicts of interest with the contents of this article.

## Abstract

Bacterial actin MreB forms filaments in which the unit of the structure is an antiparallel double strand. The wall-less helical bacterium *Spiroplasma* has five MreB homologs (MreB1–5), a part of which is composed of an intra-cellular ribbon for driving its swimming motility. The interaction modes of each ribbon component are unclear, although these are clues for understanding *Spiroplasma* swimming. Here, we examined the assembly properties of *Spiroplasma eriocheiris* MreB5 (SpeMreB5), which forms sheets and is a component protein of the ribbon. Electron microscopy (EM) revealed that sheet formation was inhibited under acidic conditions and paracrystal structures were formed under acidic and neutral conditions with low ionic strength. Solution assays found four properties of paracrystals as follows: (I) their formation followed sheet formation, (II) electrostatic interactions were required for their formation, (III) the positively charged and unstructured C-terminal region contributed to the nucleation of their formation, and (IV) their formation required Mg^2+^ at neutral pH but was inhibited by divalent cations under acidic conditions. During these studies, we found two aggregation modes of SpeMreB5, with distinct responses to ATP. These properties will shed light on SpeMreB5 assembly dynamics at the molecular level.

## Introduction

MreB belongs to the actin superfamily and is conserved in the bacterial kingdom (1). It possesses a canonical actin fold, which is composed of four subdomains (IA, IB, IIA, and IIB) (2–6). MreB molecules polymerize into antiparallel double-stranded filaments and undergo repeat polymerization and depolymerization, depending on ATP (2). The MreB filaments bind to the cell membrane via their membrane-binding sites in subdomain IA (3,7) and form an elongasome complex, which is a bacterial cell wall (peptidoglycan) synthesis complex during the growth phase (8).

Although one of the most recognized roles of MreB is elongasome formation, several bacteria use MreBs for other cellular activities (4,9–13). *Spiroplasma* belongs to the class Mollicutes and is characterized as a wall-less helical cell (14,15). Each *Spiroplasma* species possesses five classes of MreBs (MreB1–5) (16,17), and at least three of them form an intra-cellular helical ribbon structure with a *Spiroplasma* specific cytoskeletal protein fibril (18–21). The ribbon helicity is switched possibly by dynamics related to the polymerization and depolymerization of MreB2 and/or MreB5, and helicity switching is transmitted along the ribbon structure (22). These helicity-switching dynamics generate a propulsive force for a cell to swim in a liquid (23–25). This swimming is completely different from the conventional types of bacterial motility, such as flagellar and pili motility (26).

While the unit of the MreB filament is a double strand, the majority of the reported MreBs also have higher-ordered structures. Previous studies have shown that the MreBs of various species form sheet structures (2,6,27–30). In the ribbon structure of *Spiroplasma* cells, MreB(s) form sheet structures and are located between two bundles of fibril filaments and/or between a fibril bundle and the cell membrane (20,21). *Spiroplasma eriocheiris* MreB5 (SpeMreB5), an essential MreB for *Spiroplasma* swimming (4,11), forms sheet structures in vitro which include an antiparallel double-stranded filament at one edge of the sheet. The other protofilaments in the sheet are aligned parallel to adjacent protofilaments (2). In contrast, many proteins in the actin superfamily form bundles under several in vitro conditions (31). Actin formed bundles in the presence of 10–50 mM divalent cations. These bundle formations depend on the negatively charged nature of the actin filament surfaces (31–34). Several walled-bacterial MreBs have been reported to form bundles, the formation efficiencies of which depend on pH, ionic strength, and divalent cations (27–30,35). Analyses of the sheet and bundle formation processes of cytoskeletal proteins are important for understanding their behaviors at the molecular level. However, the limited experimental conditions for MreB hamper the full understanding of its sheet- and bundle-formation processes. Moreover, bundle formations of *Spiroplasma* MreBs are poorly characterized.

In this study, we analyzed the sheet and bundle formation of SpeMreB5. Electron microscopy (EM) under various conditions revealed paracrystal formation of SpeMreB5. Light scattering and sedimentation assays revealed the molecular properties of paracrystals. These findings provide clues to understand the properties of SpeMreB5. During these studies, we found two aggregation modes of SpeMreB5 that showed distinct responses to ATP.

## Results

### SpeMreB5 forms double-stranded filaments and paracrystals other than sheets

We expressed and purified monomeric SpeMreB5 by fusion with a 6×His-tag, as previously described (2). We first polymerized 10 μM SpeMreB5 by adding 2 mM Mg-ATP under three different pH conditions (50 mM CH_3_COOH-KOH pH 4.9, HEPES-KOH pH 7.0, and CHES-KOH pH 9.4 as representatives of acidic, neutral, and basic pH conditions, respectively) over a range of KCl concentrations (50, 200, and 400 mM) and observed using negative-staining EM (Fig. 1A-B, S1A-G). At pH 7 and 200 mM KCl, SpeMreB5 formed sheet structures, as observed in our previous study (Fig. S1D) (2). Sheets were also formed at pH 7 with 400 mM KCl and at pH 9 (Fig. 1B, S1E-G). Among the nine tested conditions, sheets were the dominant structure for polymerized SpeMreB5 (Fig. 1B-C). Interestingly, we found filamentous structures other than sheets under several conditions (Fig. 1C). At pH 5 and 7 with 50 mM KCl, SpeMreB5 formed paracrystal structures in which protofilaments were packed in an identical orientation (Fig. 1A, S1C). The paracrystal length reached several micrometers. These structures have also been reported in *Thermotoga maritima* MreB (TmMreB) and *Escherichia coli* MreB (EcMreB) as bundles (29,30). In contrast, at pH 5 with 200 and 400 mM KCl, SpeMreB5 formed double-stranded filaments (Fig. S1A-B), which has also been reported in *S. eriocheiris* MreB3 (SpeMreB3) and *Caulobacter crescentus* MreB (CcMreB) (2,5). We also performed the same experiments on SpeMreB3, which was purified as a monomer in our previous study (2). At pH 7, SpeMreB3 formed double-stranded filaments regardless of the KCl concentration, as in our previous study (Fig. S1H-J) (2). However, filamentous structures were not observed under the other pH conditions (Fig. S1K-L), which did not allow us to validate the bundle formations of SpeMreB3.

**Figure 1.**
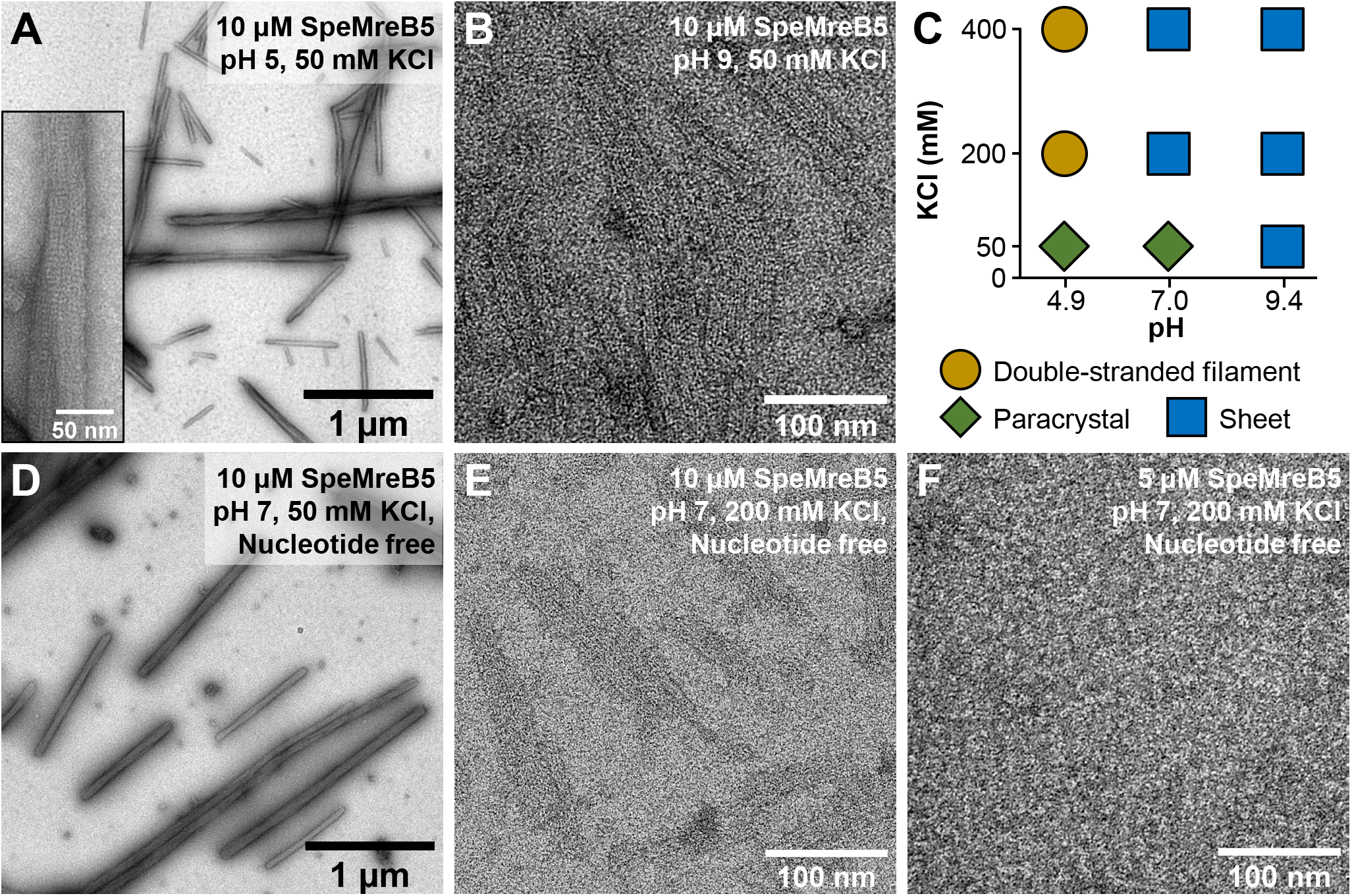
pH and ionic strength dependence of assembled SpeMreB5 structure. (**A-B**) Negative-staining EM images of 10 μM SpeMreB5 polymerized in the presence of 2 mM Mg-ATP with 50 mM KCl at pH (**A**) 5 and (**B**) 9. A magnified image of the paracrystal is shown in the inset of panel A. (**C**) A phase diagram of SpeMreB5 filament structures at 10 μM protein concentration over a range of pH and KCl concentrations. Filament structures in each condition are indicated by shapes with different colors as follows: double-stranded filament (ocher circle), sheet (blue square), and paracrystal (green diamond). (**D-F**) Negative-staining EM images of (**D-E**) 10 and (**F**) 5 μM SpeMreB5 polymerized in the absence of nucleotides at pH 7 with (**D**) 50 and (**E-F**) 200 mM KCl. Scale bars are indicated in each panel.

We also performed negative-staining EM in the absence of Mg-ATP, as a control. SpeMreB5 at pH 5 and 9, and SpeMreB3 did not form filamentous structures, as confirmed in our previous study (Fig. S1M-Q) (2). Interestingly, SpeMreB5 (at pH 7)polymerized even in the absence of Mg-ATP (Fig. 1D-E). The experimental conditions of this study differ from those of our previous study in protein concentrations (5 μM in the previous study and 10 μM in this study) and buffers (20 mM Tris-HCl pH 7.5 in a previous study and 50 mM HEPES-KOH pH 7.0 in this study) (2). In the HEPES buffer, 5 μM SpeMreB5 formed filamentous structures in the presence of Mg-ATP (Fig. S1R) but not in the absence of Mg-ATP (Fig. 1F). These results indicate that ATP promotes SpeMreB5 polymerization. In contrast, 10 μM SpeMreB5 polymerized in the absence of Mg-ATP in the Tris buffer used in the previous study (Fig. S1S) (2). These results indicate that SpeMreB5 with a concentration of approximately 10 μM or more polymerizes even without nucleotides at neutral pH.

### Surface potential maps of SpeMreB5 over the range of pH

To discuss the atomic basis of sheet and paracrystal formation, we calculated the surface potential maps of a crystal structure of the MreB5 protofilament reported previously (*Spiroplasma citri* MreB5 (SciMreB5) (PDB:7BVY) (4), which is 87.5% identical to SpeMreB5 (2)), at pH 5, 7, and 9 (Fig. 2A-C). In this study, we call the interprotofilament interaction surface for the antiparallel filament formation “back” and the opposite side “front.” We also fitted these structures to the antiparallel double-stranded filament of CcMreB (PDB:4CZJ), in which the crystal structure was reported (5), to validate potential maps of filament sides (Fig. 2D-F, S2A-C). We call the side for subdomains IA and IB “membrane side” and the opposite side “cytosolic side” as MreBs bind with the membrane via either two consecutive hydrophobic residues, an N-terminal amphipathic helix, and/or a positively charged C-terminal tail all at subdomain IA (3,7). For comparison, we calculated the surface potential maps of SpeMreB3 (PDB:7E1G), TmMreB (PDB:1JCG), and EcMreB (modeled by AlphaFold2 (36)) at pH 7, where EM studies have been conducted (2,29,30,35). Of note, we excluded the membrane sides from the following discussion, as the terminal regions of SciMreB5 and SpeMreB3 with around 20 residues that should occupy the membrane sides were not visualized (Fig. S2A-D).

**Figure 2.**
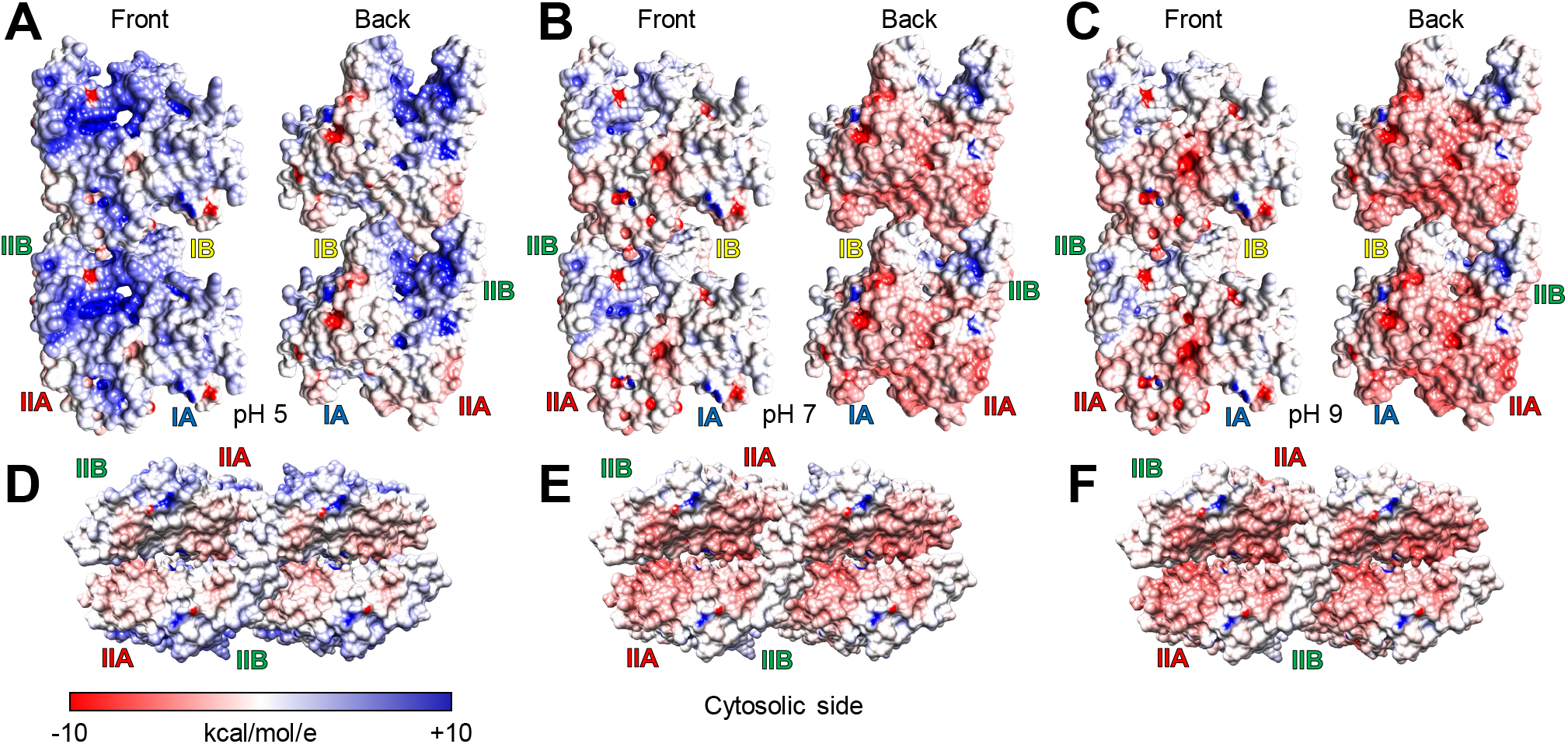
Surface potential maps of SciMreB5. The coulombic electrostatic potential is indicated by a color gradient from blue (10 kcal/mol/e) to red (−10 kcal/mol/e); namely blue, white, and red regions indicate positively charged, uncharged, and negatively charged regions, respectively. (**A-C**) Potential maps of the protofilaments with two subunits of SciMreB5 AMPPNP (PDB: 7BVY) at pH (**A**) 5, (**B**) 7, and (**C**) 9. The four subdomains are labeled for the lower subunits in the protofilament. (**D-F**) Potential maps on the cytosolic side of the double-stranded filament model of SciMreB5 AMPPNP (PDB: 7BVY) at pH (**D**) 5, (**E**) 7, and (**F**) 9. The structural model was created by fitting four SciMreB5 AMPPNP molecules to each subunit of a double-stranded filament structure of CcMreB (PDB: 4CZJ). The positions of facing subdomains (IIA and IIB) are labeled for the left side subunits.

The SciMreB5 protofilament at pH 5 was mostly positively charged on the back and front sides, whereas its cytosolic region was surrounded by weakly negatively charged regions (Fig. 2A and D). The potential map of the SciMreB5 protofilament at pH 7 differed strikingly from that at pH 5 (Fig. 2A-B and D-E). The back side of the protofilament was mostly surrounded by negatively charged regions. The charge on the front side remained weakly positive, whereas that of the subdomain IIA moiety became weakly negative (Fig. 2B). The negative charges on the cytosolic regions at pH 7 were stronger than those at pH 5 (Fig. 2D-E). These features are common to the potential maps of TmMreB and EcMreB protofilaments (Fig. S2E-F) and different from that of the SpeMreB3 protofilament, in which the front and back sides and the cytosolic region are positively charged (Fig. S2D). The overall surface charge distribution of the SciMreB5 protofilament at pH 9 was slightly different from that at pH 7 (Fig. 2B-C and E-F). A difference in the distributions is found at the subdomain IIA moiety of the front side, in which the negatively charged region becomes wider at pH 9 than at pH 7. Moreover, the negative charge on the overall structure became slightly stronger. These differences in the surface potential probably caused the various sheet and paracrystal formation modes of SpeMreB5 under various solution conditions (Fig. 1C).

### The paracrystal formations of SpeMreB5 at a neutral pH follows by disaggregation and sheet formations

To clarify the assembly dynamics of the SpeMreB5 higher-order structures observed under EM (diamonds and squares in Fig. 1C), we performed static light scattering assays. The samples were kept on ice at 5X concentrations prior to the experiments and were measured for the scattering of 650 nm light at 90° at 25 °C. In time-course measurements, we set the KCl concentration to 10 mM to promote assembly dynamics (Fig. 3A-B). At pH 7, the scattering pattern sequentially transitioned through three states over time after the addition of 2 mM Mg-ATP as follows: (I) the scattering intensity dropped to the background level in the first ~10 seconds, (II) the low scattering intensity continued for ~30 seconds as the “lag phase,” and (III) the scattering intensity dramatically increased and reached the plateau in approximately 10 min (Fig. 3A green solid line). The scattering profile at pH 5 did not have a lag phase and did not show an initial drop in intensity, unlike that at pH 7. Instead, the scattering profile showed two phases, in which the first phase reached a plateau in approximately 2 min and the second phase reached it in approximately 15 min (Fig. 3A, yellow solid line). The plateau intensity of the second phase at pH 5 was 1.6 times higher than that at pH 7, probably reflecting the difference in the heterogeneity of higher-order structures including sheets and paracrystals (also described in Fig. 6). The scattering intensity at pH 5 did not increase in the absence of Mg-ATP (Fig. 3A, yellow dotted line), indicating that Mg-ATP was a factor that increased the intensity. The initial scattering intensity at pH 7 is high in the absence of Mg-ATP. The intensity was unchanged for approximately 3 min and gradually increased to the plateau level in the presence of Mg-ATP in approximately 15 min (Fig. 3A, green dotted line).

**Figure 3.**
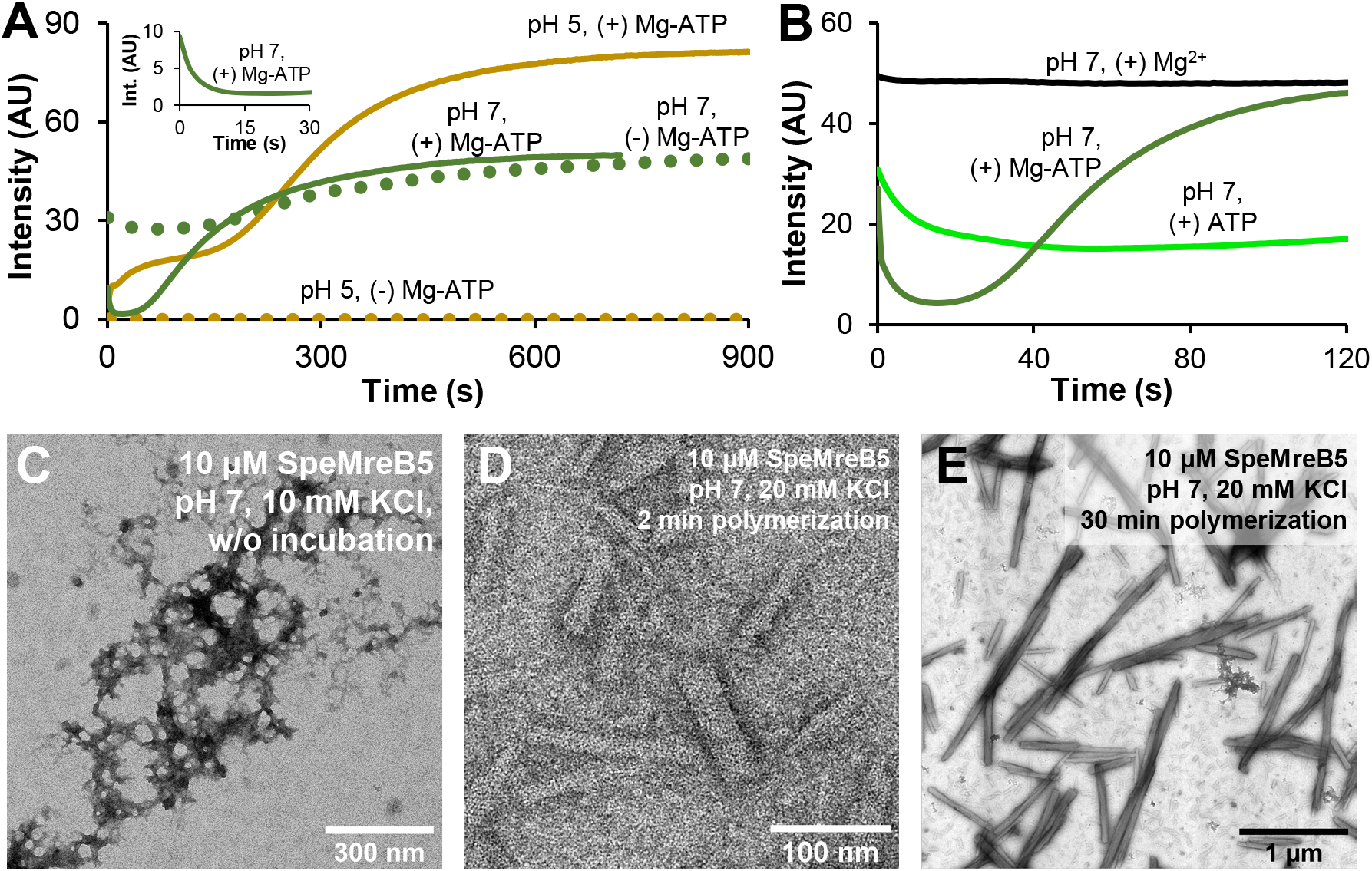
Dynamics of paracrystal formation and aggregation disassembly. For time-course light scattering, representative traces from three repeated assays for each condition are shown. (**A**) Assembly dynamics of 10 μM SpeMreB5 at pH 5 (ocher) and 7 (green) with 10 mM KCl measured by light scattering. The measurement in the presence of 2 mM Mg-ATP and in the absence of nucleotides are plotted as solid and dotted lines, respectively. The spectrum of the first 30 seconds at pH 7 with Mg-ATP is highlighted in the inset. (**B**) Assembly dynamics of 10 μM SpeMreB5 in the presence of (black) 2 mM MgCl_2_, (light green) 2 mM ATP, and (green) 2 mM Mg-ATP at pH 7 with 10 mM KCl measured by light scattering. (**C-E**) Negative-staining EM image of 10 μM SpeMreB5 (**C**) without incubation and (**D-E**) polymerized for (**D**) 2 and (**E**) 30 min at pH 7 in the presence of 2 mM Mg-ATP. KCl concentration in panel C was same as that in panels A and B (10 mM), while that in panels D and E was 20 mM to obtain the assembly dynamics with a longer lag phase than that with 10 mM KCl (see Fig. S3A). Scale bars are indicated in each panel.

To clarify the mechanism of the initial intensity drop by adding Mg-ATP at pH 7, we separately added 2 mM Mg^2+^ or 2 mM ATP to SpeMreB5 at pH 7, and observed the initial dynamics. The scattering intensity remained high after the addition of Mg^2+^ (Fig. 3B, black line). The addition of ATP alone decreased the initial scattering intensity, although it did not reach the background level (Fig. 3B, light green line), unlike the addition of Mg-ATP (Fig. 3B, green line). These results indicate that both Mg^2+^ and ATP were related to the initial intensity drop of SpeMreB5 at pH 7. In a later paragraph, we show that the initial intensity drop was induced by Ca^2+^ and ATP (Fig. S5D). To further characterize the decrease in the initial intensity, we performed the following two experiments. First, we measured the assembly dynamics of SpeMreB5 at pH 7 in the presence of 2 mM Mg-AMPPNP (an unhydrolyzed ATP analog) or Mg-ADP, instead of Mg-ATP. SpeMreB5 formed paracrystals at pH 7 in the presence of the ATP analogs (Fig. S3A-B). The plateau intensities with Mg-AMPPNP or Mg-ADP were approximately 70% compared with those with Mg-ATP, suggesting that paracrystal formation was promoted by ATP hydrolysis. Despite this difference, the scattering profiles composed of three sequential states were common among the three different nucleotide conditions (Fig. S3C). Second, we added 2 mM Mg-ATP to the sample that had already reached a plateau in the absence of nucleotides, that is, Mg-ATP addition after SpeMreB5 paracrystal formation in the absence of nucleotides (Fig. 1D). The scattering intensity remained unchanged, even after the addition of Mg-ATP (Fig. S3D), indicating that the decrease in the initial intensity of SpeMreB5 at pH 7 was caused by disruption of higher-order structures other than paracrystals by nucleotide interactions rather than hydrolysis.

To clarify the structural basis of the assembly dynamics, we observed SpeMreB5 in each state of the assembly dynamics. Prior to polymerization at pH 7, aggregating structures of sub-micrometer sizes were observed instead of filamentous structures (Fig. 3C). To visualize the lag phase in the presence of Mg-ATP at pH 7, we increased the KCl concentration to 20 mM, in which the lag phase was approximately 2 min longer than that at 10 mM KCl for ease of handling (Fig. S3E). SpeMreB5 in the lag phase formed sheet structures (Fig. 3D), together with thin and small paracrystals of sub-micrometer lengths (Fig. S3F). Paracrystal structures were formed at the plateau, as was our first EM observation (Fig. 3E, S1C). These results indicate that SpeMreB5 at neutral pH changes the assembly state in the order of aggregates, sheets, and paracrystals.

We also observed the first plateau at pH 5 in the presence of Mg-ATP using EM. However, sheet structures were not observed, unlike at pH 7. This result is consistent with our first EM observations, where sheets were not observed at pH 5 (Fig. 1C, S1A-B).

### SpeMreB5 paracrystals formations require electrostatic interactions

Our light scattering assays were able to measure SpeMreB5 paracrystal formation (Fig. 3). Using these assays, we studied the formation mechanism of SpeMreB5 paracrystals. We first measured the steady-state intensities of SpeMreB5 paracrystals over a range of ionic strengths (Fig. 4A, S3G). At both pH 5 and 7, the scattering intensities were mostly constant at 10-30 mM KCl, decreased as the KCl concentration increased at 40-70 mM KCl, and became less than the detection limit at 80 mM KCl or higher. Next, we performed disassembly assays in which pre-formed paracrystals were disrupted by changes in the solution conditions (Fig. 4B-C). The scattering intensity at both pH 5 and 7 decreased as the KCl concentration increased (Fig. 4B), indicating that the paracrystals were disrupted by increasing ionic strength. We also performed disassembly assays by changing the pH. The scattering intensities remained unchanged after the pH was shifted to 6 or 7. However, when the pH was shifted to 8 or 9, the scattering intensities decreased to the background level within 2 min (Fig. 4C). These results indicate that SpeMreB5 paracrystal formation requires electrostatic interactions.

**Figure 4.**
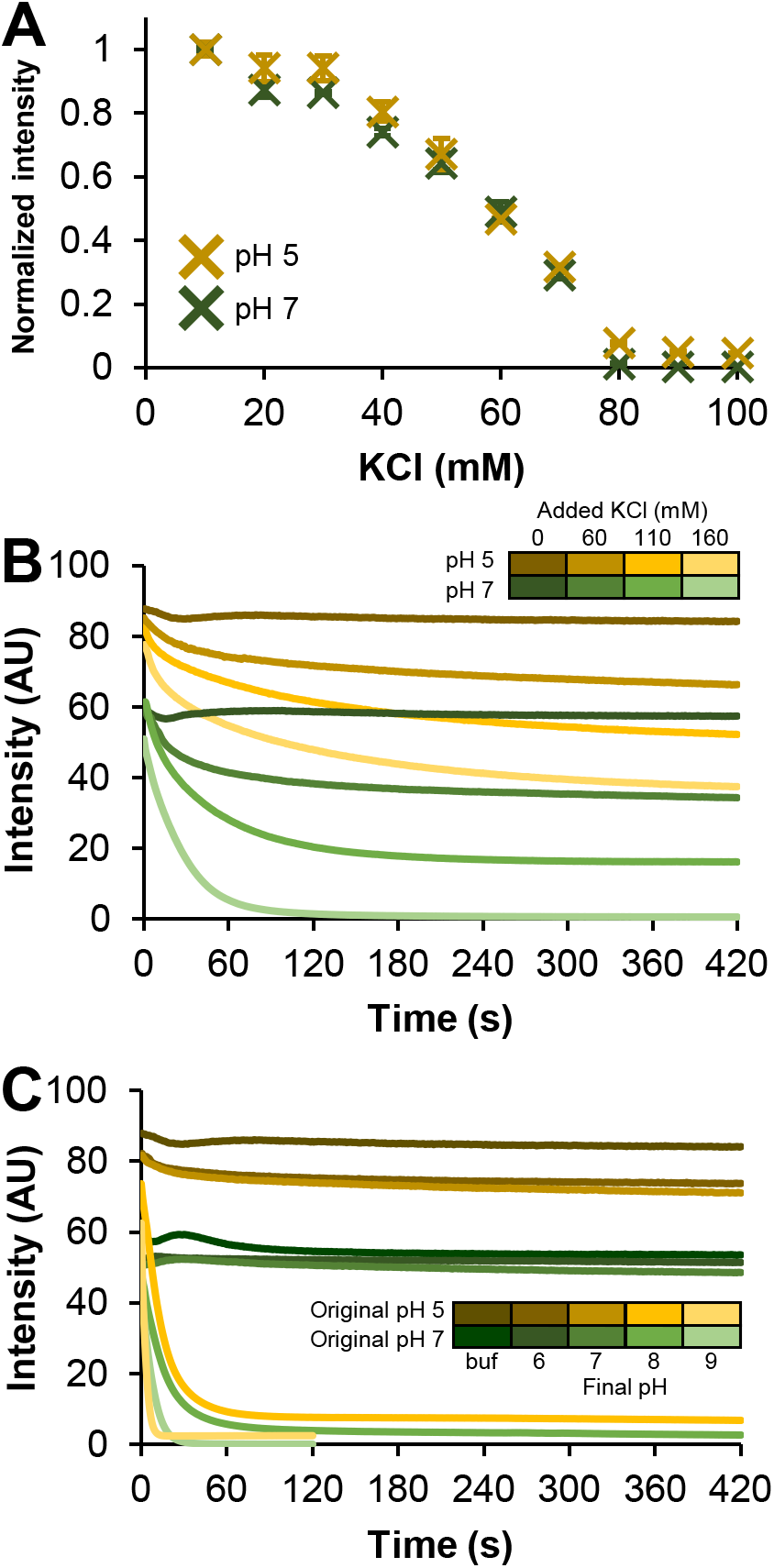
Electrostatic interaction dependence of SpeMreB5 paracrystal. For time-course light scattering, representative traces from three repeated assays for each condition are shown. (**A**) Normalized steady-state light scattering of 10 μM SpeMreB5 polymerized in the presence of 2 mM Mg-ATP over the range of KCl concentration at pH 5 (ocher) and 7 (green). Bars indicate S.D. from three independent measurements. (**B-C**) Disassembly dynamics of SpeMreB5 paracrystals measured using light scattering. The paracrystal solutions were prepared by polymerizing 10 μM SpeMreB5 with 2 mM Mg-ATP in buffers of (ocher scaled colors) 10 mM CH_3_COOH-KOH pH 4.9 and (green scaled colors) 10 mM HEPES-KOH pH 7.0 with 40 mM KCl. The measurements in which the buffer composition was unchanged are indicated with the thickest colored lines. (**B**) Disassembly was induced by increasing KCl concentration into (second thickest colored line in each color scale) 100, (second thinnest colored) 150, and (thinnest colored) 200 mM. (**C**) Disassembly was induced by changing the buffer pH into (second thickest colored line in each color scale) 6, (medium colored) 7, (second thinnest colored) 8, and (thinnest colored) 9 by adding 50 mM MES-KOH pH 6.0, HEPES-KOH pH 7.0, HEPES-KOH pH 8.1, and CHES-KOH pH 9.4, respectively.

### The positively charged C-terminal region of SpeMreB5 is contributed to the nucleation of paracrystals and the aggregation at a neutral pH

Each *Spiroplasma* MreB5 possesses a C-terminal unstructured region with a positive net charge (3,4,16). To investigate the effects of this region on paracrystal formation, we prepared two SpeMreB5 variants with truncations of 9 and 26 residues at the C-terminus (ΔC9 and ΔC26, respectively) (Fig. 5A). SpeMreB5 ΔC9 is comparable to SciMreB5 ΔC10 used in a previous study (3), and SpeMreB5 ΔC26 is a variant in which all C-terminal residues outside the MreB folding domain are removed. Thus, ΔC9 was successfully purified. In contrast, ΔC26 did not bind to the Ni^2+^-NTA affinity column, possibly due to non-specific binding between the 6×His-tag and the ΔC26 surface. Moreover, the solubility of ΔC9 at pH 5 was not high enough for polymerization experiments, although the solubility at pH 7 was sufficient. Therefore, we analyzed the paracrystal formation of the C-terminal-truncated variant of SpeMerB5 using ΔC9 at pH 7 by light scattering. The initial intensity of ΔC9 was the background level, unlike the wild-type (WT) (Fig. 3A, 5B), indicating that the initial aggregation of SpeMreB5 (Fig. 3C), which caused the high initial intensity, was formed via the C-terminal region. ΔC9 in the presence of Mg-ATP assembled slower than that of WT and did not reach a plateau after 20 min (Fig. 5B). However, the steady-state intensities of ΔC9 were slightly lower than those of WT over the KCl concentrations (Fig. 5C). To evaluate whether the slow assembly of ΔC9 was caused by the sheet assembly phase, we performed sedimentation assays under the conditions used in our previous study in which SpeMreB5 formed sheets but did not form paracrystals (2). Using these assays, we estimated the critical concentrations that reflect the minimum concentration required for polymerization and the ratio of the dissociation and association rates (Fig. S4A-B and E). Those of WT and ΔC9 were not significantly different (Table 1), suggesting that truncation of the C-terminal 9 residues did not affect the dynamics of polymerization and sheet formation. These results indicated that the C-terminal region of SpeMreB5 is involved in the nucleation step of paracrystal formation.

**Figure 5.**
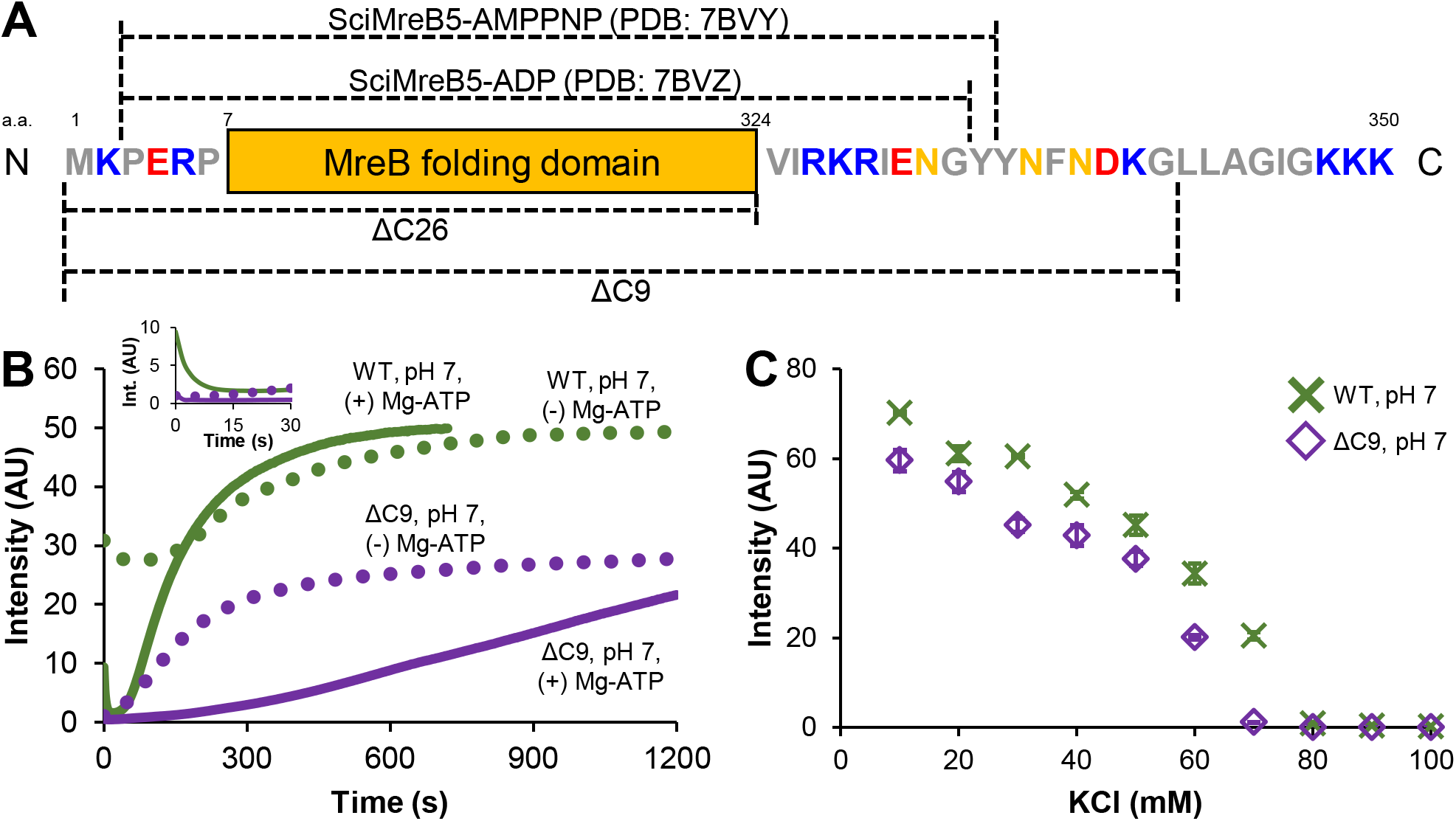
Assembly dynamics of paracrystals by the C-terminus truncated variant of SpeMreB5. (**A**) Schematics of the SpeMreB5 sequence. The MreB folding domain is defined as the visible region in all the MreB crystal structures reported previously (2–6) and invisible-flexible loops within them. The residues exterior of the MreB folding domain are shown with gray, orange, red, and blue colors for hydrophobic, non-polar-hydrophilic, acidic, and basic ones, respectively. The regions that are visible in the previously reported crystal structures of SciMreB5 (3,4) and for SpeMreB5 ΔC9 and ΔC26 variants are indicated above and underneath the schematics, respectively. (**B**) Assembly dynamics of 10 μM SpeMreB5 WT (green, the same traces as those in Fig. 3A) and ΔC9 variant (purple) at pH 7 with 10 mM KCl measured using light scattering. The measurement in the presence of 2 mM Mg-ATP and in the absence of nucleotides are plotted as solid and dotted lines, respectively. Representative traces from three repeated assays for each condition are shown in the graph. The spectrums of the first 30 seconds are highlighted in the inset. (**C**) Steady-state light scattering of 10 μM SpeMreB5 WT (green, the same plots as those in Fig. S3B) and ΔC9 (purple) polymerized in the presence of 2 mM Mg-ATP over the range of KCl concentration at pH 7. Bars indicate S.D. from three independent measurements.

**Table 1.** Bulk critical concentrations of SpeMreB5 WT with varying divalent cation conditions and ΔC9 variant measured using a sedimentation assay.^a^.

| | 2 mM Mg-ATP<br>WT | 2 mM Mg-ATP<br>ΔC9 | 2 mM Ca-ATP<br>WT | 2 mM ATP<br>(X<sup>2+</sup> free) WT |
| --- | --- | --- | --- | --- |
| Cc of SpeMreB5<br>(μM) | 0.09 ± 0.01 | 0.11 ± 0.04 | 0.09 ± 0.08 | 0.05 ± 0.03 |
| *p*-value v.s. 2 mM<br>Mg-ATP WT | - | 0.45 | 0.95 | 0.07 |

<sup>a</sup>. The values are indicated as mean ± S.D. from three repeated measurements. The row below the critical concentrations indicates the *p*-value against the critical concentration in the presence of 2 mM Mg-ATP supported by Student's *t*-test, showing that the significance of critical concentration differences is not supported for all tested pairs.

In the absence of Mg-ATP, the scattering intensity of ΔC9 increased immediately after initiating the measurement and plateaued at the same timescale as the WT in the absence of Mg-ATP. However, the plateau intensity of ΔC9 in the absence of Mg-ATP was half that of WT in the absence of Mg-ATP (Fig. 5B). ΔC9 at this condition formed paracrystals with lengths shorter than 1 μm (Fig. S1T). These results suggest three possibilities: (I) the reaction path of WT and ΔC9 in the absence of Mg-ATP is identical; (II) SpeMreB5 undergoes different reaction paths with and without nucleotides, and (III) paracrystal formation in the absence of Mg-ATP is promoted by aggregate formation.

### Paracrystal formations of SpeMreB5 require Mg^2+^ at a neutral pH but are inhibited by divalent cations at an acidic pH

Previous studies have revealed that bundle formation of actin superfamily proteins requires divalent cations (27,28,31,32). We then examined the requirement of divalent cations for SpeMreB5 polymerization and the formation of higher order structures. First, we performed sedimentation assays on SpeMreB5 under various divalent cation conditions (Fig. S4). The pellet amounts of SpeMreB5 were mostly constant over Mg^2+^ and Ca^2+^ concentrations (Fig. S4F-H). The critical concentrations of SpeMreB5 were not significantly different among conditions in the presence of 2 mM ATP (divalent cation-free), Mg-ATP, and Ca-ATP (Fig. S4A and C-E, Table 1). These results indicate that SpeMreB5 does not require divalent cations for polymerization or sheet formation.

We also examined the effects of divalent cations on SpeMreB5 paracrystal formations. At pH 7, in the presence of 2 mM Ca-ATP, SpeMreB5 formed sheets, while the corresponding condition in the presence of Mg-ATP formed paracrystals (Fig. 6A, S1C). In contrast, paracrystals were formed at pH 5 in the presence of 2 mM Ca-ATP (Fig. S5A) as well as in the presence of Mg-ATP (Fig. 1A). In the divalent cation-free condition at pH 7, most of SpeMreB5 formed sheets (Fig. 6B) and a few fractions formed paracrystals (Fig. S5B). Surprisingly, under divalent cation-free conditions at pH 5, amorphous aggregates were observed instead of filamentous structures (Fig. 6C).

**Figure 6.**
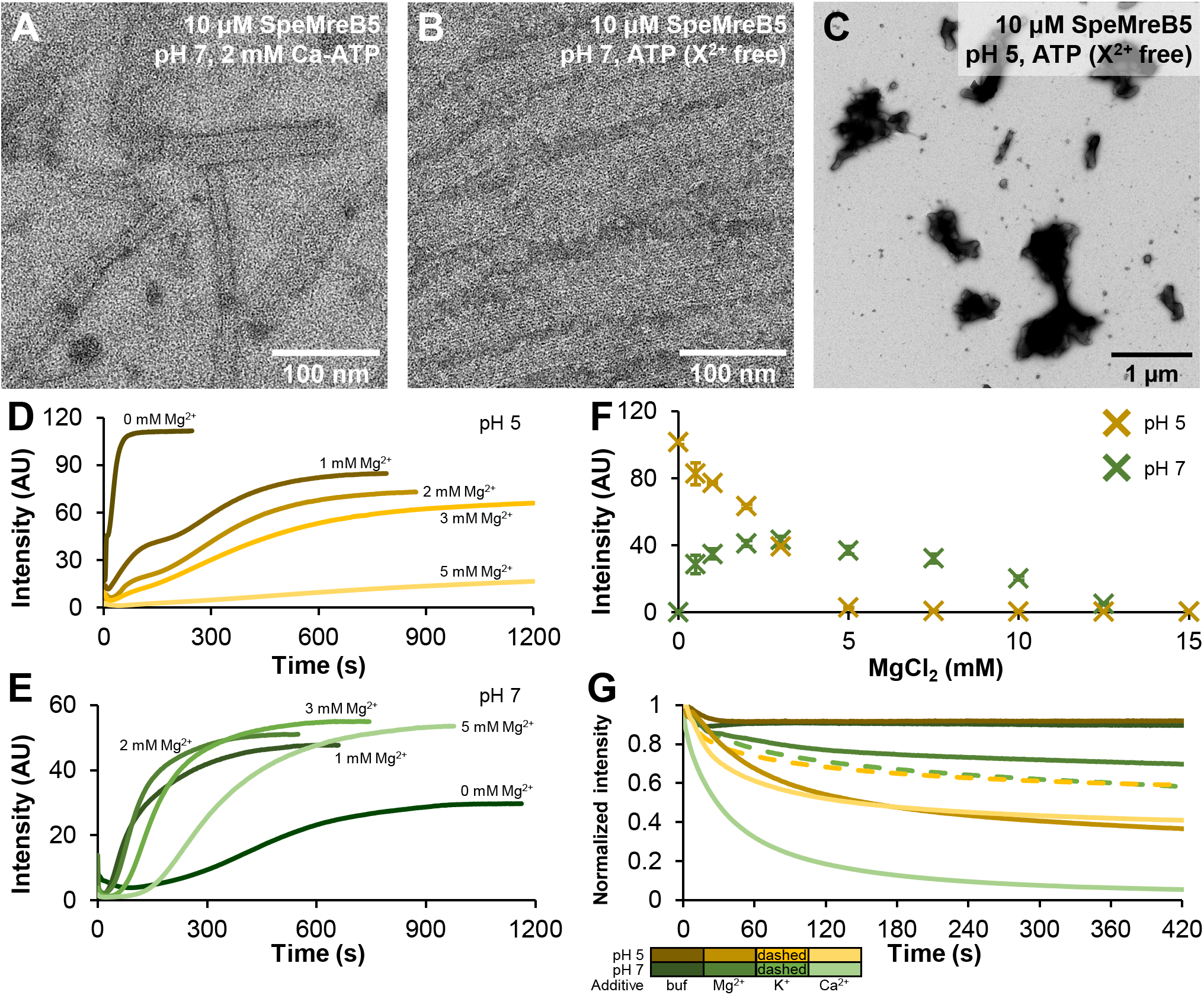
Divalent cation dependence of SpeMreB5 polymerizations and higher-order structure formations. For time-course light scattering, representative traces from three repeated assays for each condition are shown. For divalent cation-free conditions, 1 mM EDTA-NaOH pH 8.0 was added to avoid effects from contaminating amounts of multivalent cations. (**A-C**) Negative-staining EM images of 10 μM SpeMreB5 polymerized with varying divalent cation conditions with 50 mM KCl. SpeMreB5 was incubated at pH (**A-B**) 7 and (**C**) 5 in the presence of (**A**) 2 mM Ca-ATP and (**B-C**) 2 mM ATP (divalent cation-free). Scale bars are indicated in each panel. (**D-E**) Mg^2+^-dependent assembly dynamics of 10 μM SpeMreB5 at pH (**D**) 5 and (**E**) 7 with 10 mM KCl measured using light scattering. The polymerization was initiated by adding 2 mM ATP with varying MgCl_2_ concentrations as indicated in the color scales in the panels. (**F**) Steady-state light scattering of 10 μM SpeMreB5 polymerized with 2 mM ATP at pH (ocher) 5 and (green) 7 over the range of MgCl_2_ concentration. KCl concentration was 50 mM constant. Bars indicate S.D. from three independent measurements. (**G**) Normalized light scattering traces of disassembly dynamics of SpeMreB5 paracrystals induced with divalent cations. The paracrystal solutions were prepared by polymerizing 10 μM SpeMreB5 in the presence of 2 mM Mg-ATP at pH (ocher scaled colors) 5 and (green scaled colors) 7 with 50 mM KCl. Disassembly was induced by adding (second thickest colored solid line) 15 mM MgCl_2_, (thinnest colored solid line) 15 mM CaCl_2_, and (second thinnest colored dashed line) 60 mM KCl. Of note, the ionic strengths of these salts are identical assuming the same degree of dissociation (see Materials and Methods). The measurements in which the buffer condition was unchanged are indicated with the thickest colored solid lines.

To clarify the effects of divalent cations on the assembly dynamics of paracrystals, we performed time-course measurements of SpeMreB5 paracrystal assembly (Fig. 6D-E, S5C-D). In the divalent cation-free condition at pH 5, the scattering profile was single-phase and plateaued at the same time scale as the first phase in the presence of Mg^2+^, indicating that the first phase of assembly dynamics at pH 5 reflects the aggregation of SpeMreB5 (Fig. 6C-D). The plateau intensities of both the first and second phases in the presence of Mg^2+^ at pH 5 decreased as Mg^2+^ concentration increased. In particular, the first phase was indistinguishable in the presence of 3 mM and 5 mM Mg^2+^. Moreover, the assembly rate of the second phase decreased in an Mg^2+^-dependent manner (Fig. 6D). Ca^2+^ showed similar effects to Mg^2+^ on SpeMreB5 paracrystal formation at pH 5, while its inhibition efficiencies were less than that of Mg^2+^ (Fig. S5C). These results indicate that SpeMreB5 paracrystal formation at pH 5 is inhibited by divalent cations.

In contrast, the assembly dynamics of paracrystals at pH 7 showed puzzling Mg^2+^ dependence. Under divalent cation-free conditions, the lag phase continued for approximately 3 min, and the intensity reached a plateau in approximately 20 min. Notably, as we performed the time-course measurements under a KCl concentration of 10 mM, which is lower than the EM observations, the scattering intensity was high even though there were few paracrystal structures under the EM observation (Fig. 6E, S5B). In the presence of 1–3 mM Mg^2+^, the times for the lag phase and reaching the plateau were five and two times shorter, respectively, and the plateau intensities were two times higher than those in the divalent cation-free condition. The plateau intensity in the presence of 5 mM Mg^2+^ was not different from those in the presence of 1–3 mM Mg^2+^, whereas the lag phase became slightly longer (Fig. 6E). We also examined the effects of Ca^2+^ on paracrystal assembly at pH 7. The initial intensity rapidly decreased with the addition of 2 mM Ca-ATP, indicating that the initial aggregation (Fig. 3C) was also dissociated by Ca-ATP. Paracrystal assembly was suppressed in the presence of Ca^2+^. In particular, the intensity did not substantially increase within 20 min in the presence of 5 mM Ca^2+^ (Fig. S5D). These results indicate that SpeMreB5 paracrystal formations at pH 7 requires Mg^2+^.

Next, we measured steady-state intensities over a range of Mg^2+^ concentrations (Fig. 6F). We set the KCl concentration to 50 mM for consistency with EM observations. At pH 5, the scattering intensities decreased in a Mg^2+^-dependent manner and reached the detection limit at Mg^2+^ concentrations of 5 mM or higher (Fig. 6F, yellow). Of note, this result may overrate the Mg^2+^ effects on paracrystals, as the scattering intensities are likely derived from both paracrystal and aggregated structures (Fig. 1A, 6C). However, it is plausible that both aggregation and paracrystal formation were inhibited by Mg^2+^ at pH 5. At pH 7, the scattering intensities peaked in the presence of 3 mM Mg^2+^ and were nearly the background level at 0 and 15 mM Mg^2+^ (Fig. 6F, green), suggesting that the paracrystal formation efficiency at pH 7 is determined by the balance of the Mg^2+^ requirement for paracrystal formation and ionic strength effects. We also evaluated the paracrystal disassembly by increasing the concentration of divalent cations (Fig. 6G, S5E). At pH 5, the decrease in the scattering intensity upon the addition of MgCl_2_ and CaCl_2_ was greater than that by KCl with the same ionic strength (when the degree of dissociation is assumed to be 1 for all salts) (yellow lines in Fig. 6G and S5E). This phenomenon is common for CaCl_2_ addition at a pH of 7. On the other hand, the decrease in the scattering intensity by MgCl_2_ at pH 7 was less than that by KCl (green lines in Fig. 6G and S5E), consistent with the Mg^2+^ requirement for paracrystal formation at pH 7 (Fig. 6E-F). Disassembly assays were also performed by adding EDTA (a chelating agent for multivalent cations) at pH 7. However, the decrease in the scattering intensity by EDTA was only slightly different from that of a buffer with a pH identical to that of the EDTA solution (Fig. S5F), suggesting that Mg^2+^ was occluded in the paracrystals and was not accessed by the EDTA. Altogether, our results demonstrate that SpeMreB5 paracrystal formation at pH 5 is inhibited by divalent cations, but that at pH 7 requires Mg^2+^.

## Discussion

In this study, we investigated sheet and paracrystal formation of SpeMreB5 using EM and bulk biochemical assays. The transitions of the six states were observed in this study (Fig. 7). We discuss the properties of each state and their possible roles in *Spiroplasma* swimming in some states.

**Figure 7.**
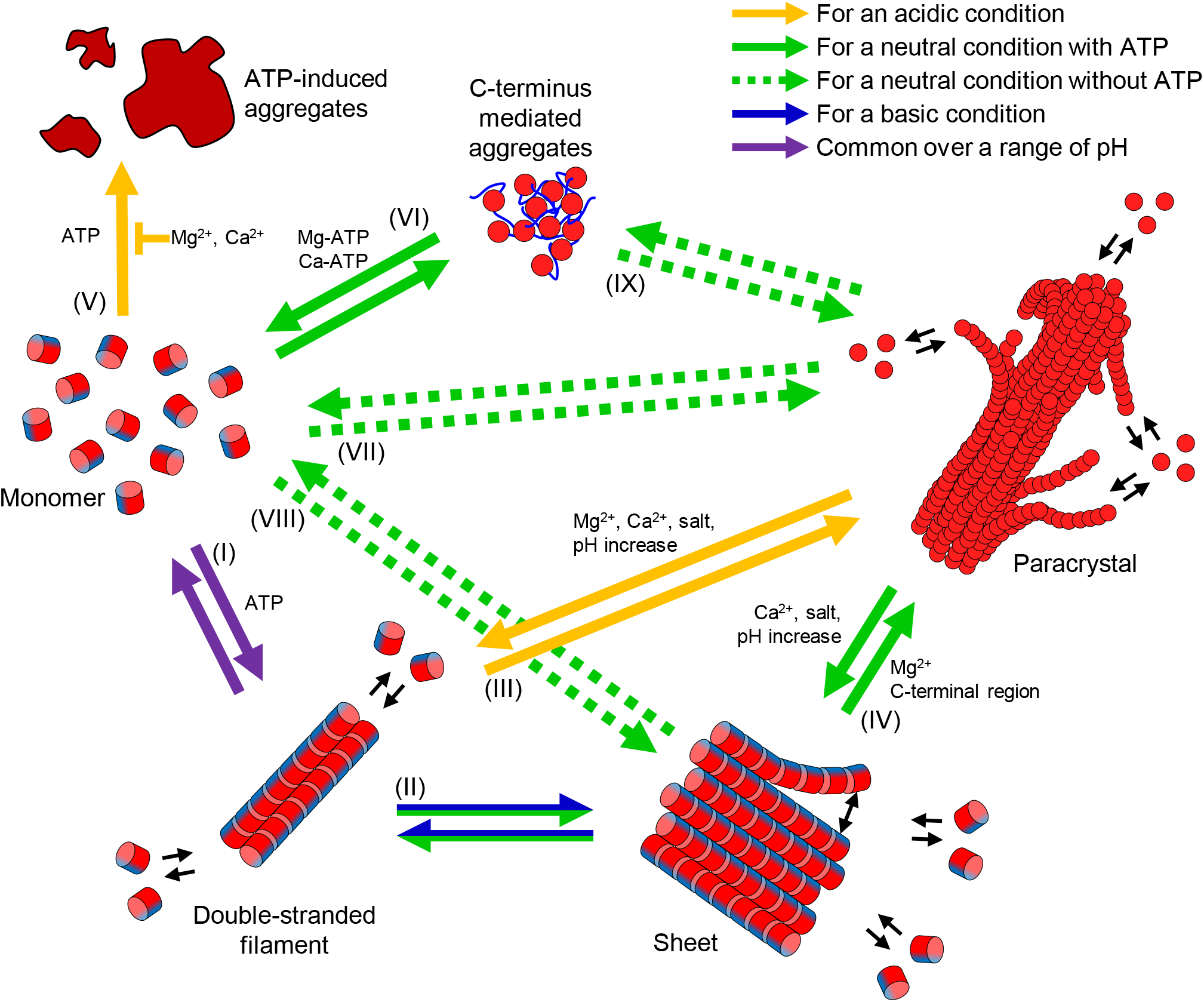
Summary for SpeMreB5 polymerization. The relationship among the six states found in this study (monomer, double-stranded filament, sheet, paracrystal, C-terminus mediated aggregates disassembled by ATP, and aggregates induced by ATP) is suggested. An SpeMreB5 subunit is indicated by a red circle or a cylinder colored with red and blue. The positively charged-unstructured C-terminus is shown as a blue line for the state of C-terminus mediated aggregates. Reactions specific for acidic, neutral, and basic pH conditions in the presence of nucleotides are shown in solid arrows colored with orange, green, and blue, respectively. Reactions common over a range of pH are indicated using purple solid arrows. Reactions specific for conditions in the absence of nucleotides are shown in green dotted lined arrows. Factors that promote a reaction step are shown alongside each arrow. An inhibition factor against a reaction step is indicated by T.

### Polymerization and sheet formations

ATP was required for polymerization at a low SpeMreB5 concentration and at acidic and basic pH, although SpeMreB5 at pH 7 formed filamentous structures under nucleotide-free conditions (Fig. 1). These results indicate that ATP promotes SpeMreB5 polymerization (Fig. 7-(I)), confirming our previous finding that nucleotide binding induces polymerization of SpeMreB3 and SpeMreB5 (2). SpeMreB5 formed double-stranded filaments at acidic pH, and sheets at neutral and basic pH (Fig. 1C). Our previous study revealed that SpeMreB5 sheets are composed of an antiparallel double-stranded filament at one edge and a parallel alignment of protofilaments (2). The minimum unit of SpeMreB5 filamentous structures that we found was the double strand as well as the findings of a previous study (Fig. 1C, S1A-B) (3), suggesting that sheet formation likely initiates from the double-stranded filaments (Fig. 7-(II)). The SciMreB5 protofilament is mostly surrounded by positively charged regions on both the front and back sides at pH 5, while the back side is negatively charged at pH 7 and 9 (Fig. 2A), suggesting that sheet formation at an acidic pH is inhibited by electrostatic repulsion between the front and back sides. This is consistent with the results, in which SpeMreB3 at pH 7 did not form sheets (Fig. S1H-I), and is surrounded by positively charged regions on both the front and back sides (Fig. S2D). Considering the protofilament orientations within the sheets, the same protofilament side (Fig. 2D-F) faces the cytosolic region. This alignment indicates that SpeMreB5 sheets face wide negatively charged surfaces in the cytoplasm when they bind to the membrane. A previous study reported that SciMreB5 interacts with fibril (4). Assuming that the MreB sheet between a fibril bundle and the membrane is composed of MreB5 (20), it probably interacts with fibril filaments via the wide negatively charged regions on the cytosolic side of the sheets (Fig. 2D-F).

### Paracrystal formations

SpeMreB5 formed paracrystals under low-ionic-strength conditions at acidic and neutral pH (diamonds in Fig. 1C). While this structure has not been observed in *Spiroplasma* cells (20,21), it possibly reflects the properties of SpeMreB5 because there are clear differences in the tendency of paracrystal formation for SpeMreB3 and SpeMreB5 (Fig. 1C, S1H-J). The paracrystal formation mechanisms at acidic and neutral pH are likely different, as the paracrystals showed distinct divalent cation dependences at these two pH conditions (Fig. 6D-G). Under acidic conditions, in which sheet formation was inhibited (Fig. 1C), paracrystals likely grew from double-stranded filaments as nuclei (Fig. 7-(III)). The surface potential map of the SciMreB5 protofilament at pH 5 showed that the cytosolic side was negatively charged, and the front side, which was exposed to the solvent when the double-stranded filaments were formed, was positively charged (Fig. 2A and D), suggesting that these sides interact with each other to form paracrystals. The lengths of the paracrystals reached several micrometer orders, while those of the double-stranded filaments were sub-micrometer orders (Fig. 1A, S1A-B), indicating that paracrystal formations stabilize each protofilament.

In contrast, SpeMreB5 formed sheets at neutral pH prior to paracrystal formation in the presence of Mg-ATP (Fig. 3D-E, S3F), suggesting that the sheets work as nuclei of paracrystals (Fig. 7-(IV)). As the steady-state intensities of ΔC9 were only slightly different from those of the WT (Fig. 5C), the C-terminal region was probably not involved, and the negatively charged cytosolic region (Fig. 2E) was likely involved in the interactions for paracrystal formation. The positively charged front side was the sole candidate interaction partner for the negatively charged cytosolic region (Fig. 2B). The dependence of the front side on paracrystal formation is also supported by the inability of paracrystal formation at pH 9. The negatively charged region of the subdomain IIA moiety at pH 9 became wider than that at pH 7 (Fig. 2B-C), possibly leading to electrostatic repulsion for the inhibition of paracrystal formation. Although the front side is exposed to the solvent when SpeMreB5 forms sheets, it is unreasonable that the paracrystals are formed by sheet stacking, considering the tight packing of protofilaments within the paracrystals (Fig. 1A). Instead, it is most plausible that single protofilaments elongate on negatively charged surfaces, such as the cytosolic side of the sheets, facing the front side to grow into paracrystals. This model is consistent with previous findings in which SpeMreB5 and SciMreB5 not only form inter-protofilament interactions for antiparallel double-stranded filaments (2–4). Although the C-terminal unstructured region is expected not to be involved in interactions for paracrystal formation (Fig. 5C), this region was involved in paracrystal nucleation (Fig. 5B), suggesting that non-specific interactions via the C-terminal region increase the local concentration of SpeMreB5 around sheets and paracrystals to promote their assemblies. This idea is consistent with a previous study in which engineered proteins with flexible tubulin-binding regions on the outside of microtubules induced supra-structural formations such as microtubule doublets and branched microtubules (37). As the C-terminal region of SciMreB5 is involved in binding to the negatively charged *Spiroplasma* membrane (3), the paracrystal nucleation property may induce MreB5 nucleation on the membrane. Paracrystal formation at a neutral pH required Mg^2+^ and was inhibited by Ca^2+^ (Fig. 6E-F, S5D), suggesting that SpeMreB5 has binding sites specific for Mg^2+^. These regions probably localize on the surface of the SpeMreB5 protofilaments as those for the bundle formation of actin filaments are on its surface (32). Inhibition of bundle formation by Ca^2+^ has also been reported for EcMreB (29), suggesting that Mg^2+^ binding sites for paracrystal formation are negatively charged regions common between SpeMreB5 and EcMreB, such as the moieties of subdomains IIA and IIB on the back side of the protofilament. Paracrystal formation was inhibited by divalent cations at acidic pH (Fig. 6D, S5C), suggesting that the putative Mg^2+^ binding regions at neutral pH turn their charges by pH shifts between 5 and 7.

Bundle formation dependent on divalent cations has also been reported for actin filaments. However, the optimal concentration of divalent cations for actin bundle formations is 10–50 mM which is approximately 10 times higher than the optimal concentration for SpeMreB5 paracrystal formations (1-5 mM) (Fig. 6F) (32). Actin bundles are formed by bridging divalent cations with nine acidic residues (32), suggesting that electrostatic interactions are less involved in actin bundle formation than in SpeMreB5 paracrystal formation. This likely explains why actin bundles are resistant to the presence of high concentrations of divalent cations, which are high enough to disrupt SpeMreB5 paracrystals (Fig. 6F). Moreover, actin filaments adopt a right-handed helix, which can restrict inter-filament interactions to form bundles. On the other hand, the SpeMreB5 sheets are not helical (Fig. 1B and E, S1D-G), suggesting that small amounts of Mg^2+^ will affect paracrystal formation, as inter-protofilament interactions for paracrystal formation are unlikely to be restricted.

### Aggregations induced and disassembled by ATP

We also found two SpeMreB5 aggregates that showed distinct responses to ATP (Fig. 3C, 5C). One was formed in the presence of ATP at acidic pH (Fig. 6C-D, 7-(V), S5C). This aggregation is surprising because, to the best of our knowledge, aggregations dependent on ATP have not been reported for the actin superfamily proteins. The other was formed at a neutral pH via the C-terminal region (Fig. 3C, 5B). This aggregation was disassembled by ATP, and the disassembly efficiency increased in the presence of divalent cations (Fig. 3A-B, 7-(VI), S5D). We cannot rule out the possibility that this aggregation was affected by the initial conditions of our experiments (5X concentration prior to the assays, also described in Fig. 3). However, disassembly of aggregates by ATP is intriguing because, to the best of our knowledge, a comparable phenomenon has not been reported for actin. Although only the nucleotide-binding pocket has been reported for the ATP-binding site of MreBs (2–6), this site is unlikely to dominate the disaggregation phenomenon. ATP is not only known as a molecular unit of currency in life but is also reported as a hydrotrope of proteins (38). ATP as the hydrotrope binds nonspecifically to the termini and loops of proteins (39). This is likely the case for SpeMreB5. The increased efficiency of SpeMreB5 disaggregation in the presence of divalent cations (Fig. 3B, S5C) suggests that ATP binding for disaggregation is promoted by the coordination of a divalent cation to the tri-phosphate group of ATP. In addition, the disaggregation by ATP of SpeMreB5 is likely distinct from that of the FUS protein, which does not rely on divalent cations (38).

### SpeMreB5 polymerization in the absence of nucleotides

We found that SpeMreB5 polymerized in the absence of nucleotides at a neutral pH when the protein concentration was sufficiently high (Fig. 1D-F, S1S). This finding is intriguing because the polymerization of SpeMreB5 and SciMreB5 is thought to require ATP (2–4). The lag phase of ΔC9 in the nucleotide-free condition was indistinguishable, unlike that in the presence of Mg-ATP (Fig. 5B), suggesting that SpeMreB5 under nucleotide-free conditions undergoes reaction paths to directly shift the states from monomer to a structural state in the steady state (Fig. 7-(VII) and (VIII)). SpeMreB5 WT, which forms aggregates disassembled by ATP (Fig. 3C), in the absence of nucleotides reached a plateau intensity higher than that at ΔC9 without any intensity drops (Fig. 5B), suggesting the following two possibilities: there is a reaction path from the aggregates to paracrystals (Fig. 7-(IX)) and the local concentration increase by the aggregation facilitates paracrystal formation in the absence of nucleotides.

## Conclusions

In this study, we clarified the interactions in SpeMreB5 sheets and the formation mechanisms of paracrystals followed by sheet formation (Fig. 7) by focusing on ionic strength and pH dependence (Fig. 4), nucleation by the C-terminal region (Fig. 5), and distinct divalent cation dependences (Fig. 6). These findings will aid in the understanding of the molecular properties of SpeMreB5. In this study, we found two aggregation modes of SpeMreB5, with distinct responses to ATP (Fig. 3C, 6C). This finding also sheds light on the protein aggregation phenomena.

## Materials and Methods

### SpeMreBs expression and purification

SpeMreB3 and SpeMreB5 and its ΔC variants were expressed and purified as previously described (2). In brief, SpeMreB3 or SpeMreB5 were expressed in *E. coli* BL21 (DE3) or C43 (DE3) carrying *spemreB3* or *spemreB5* fused pCold-15b for 24 h at 15 °C with induction conditions of OD_600_ = 0.4 and IPTG = 1 mM. Cell pellets from 1 L of culture were resuspended in 20-40 mL of buffer A (50 mM Tris-HCl pH 8.0, 300 mM NaCl, 50 mM imidazole-HCl pH 8.0), sonicated, centrifuged, and purified using a HisTrap HP column (Cytiva, Wauwatosa, WI) and a HiLoad 26/600 Superdex 200 pg. column (Cytiva) equilibrated with buffer B (20 mM Tris-HCl pH 8.0, 300 mM NaCl) at 4 °C.

### SpeMreB polymerization

For the sedimentation assay, SpeMreBs were polymerized with buffer S (20 mM Tris-HCl pH 8.0, 1 M NaCl, 200 mM L-Arginine-HCl pH 8.0, 5 mM DTT, 2 mM MgCl_2_, and 2 mM ATP) (2) which inhibited polymerization and amorphous aggregation under nucleotide-free conditions. The solution conditions for other experiments are shown in the figure legend. The pH 5, 7, and 9 conditions were mediated by 50 mM CH_3_COOH-KOH pH 4.9, HEPES-KOH pH 7.0, and CHES-KOH pH 9.4, respectively, unless otherwise stated. All samples tested in this study contained 5 mM DTT to prevent unfavorable oligomerization of SpeMreBs via the S-S bonds. Prior to polymerization, the SpeMreB buffer was exchanged from buffer B to the desired buffer in the absence of DTT, divalent cation salts, and ATP. SpeMreBs with a concentration lower than the desired concentration were concentrated using an Amicon Ultra 10 K (Merck). Samples with the desired SpeMreB concentration were centrifuged (20,000×*g* at 4°C for 10 min) to remove aggregates and added for DTT, MgCl_2_, and ATP to initiate polymerization. For nucleotide-free conditions, divalent cation salts and ATP were excluded from the buffer.

### Electron microscopy (EM)

As previously described (2), a sample (4 μL) was placed onto a 400-mesh copper grid coated with carbon for 1 min at room temperature (24-27°C), washed with 10 μL water, stained for 45 s with 2% (w/v) uranyl acetate, air-dried, and observed under a JEOL JEM-1010 transmission electron microscope (Akishima, Japan) at 80 kV, equipped with a FastScan-F214T CCD camera (TVIPS, Gauting, Germany).

### Estimation of surface potential map

The surface potential maps were calculated with the PDB2PQR (40,41) server using the PARSE forcefield (42,43) in conjunction with PROPKA (44–46) to assign the protonation state at the provided pH conditions (47). Calculations were performed for each subunit, excluding the bound ligands. For the calculation of SpeMreB3 (PDB:7E1G), in which lysine residues were di-methylated prior to crystallization (2), the di-methylated lysine residues were replaced with the most probable rotamers of lysine in Dunbrank’s rotamer library (48) because the positive charge of a di-methylated lysine residue is weaker than that of un-methylated lysine. The surface potentials were visualized using Chimera ver 1.13.1 (49) with a color gradient from −10 kcal/mol/e (red) to +10 kcal/mol/e (blue).

### Sedimentation assay

SpeMreBs in buffer S polymerized at room temperature with a volume of 200 μL were polymerized for 1–6 h and centrifuged (100,000 rpm at 23 °C for 120 min) in a TLA-100 rotor (Beckman Coulter) (2). The pellet was resuspended in 200 μL water. The supernatant and pellet fractions were subjected to electrophoresis on a 12.5% Laemmli gel and stained with Coomassie Brilliant Blue R-250 to analyze the concentration of each fraction. The band intensities of SpeMreBs were quantified using ImageJ software (National Institutes of Health; http://rsb.info.nih.gov/ij/). The concentrations of the supernatant and pellet fractions were estimated as the products of the total SpeMreB concentration and the ratio of each fraction to the sum of the supernatant and pellet fractions.

### Static light scattering

Ninety-degree perpendicular light scattering experiments were carried out using FP-6200 (JASCO, Tokyo, Japan) in a single cuvette containing 60 μL sample solution at 25 °C under the control of a temperature stabilizer. Both the excitation and emission wavelengths were set to 650 nm. For the time-course measurements, a sample with protein and buffer concentrations five times higher than the desired was kept on ice prior to the assay, diluted in a solution at room temperature to mediate the composition, and immediately applied to the measurements with a lag time of approximately 5 s due to manual mixing. For steady-state light scattering measurements, SpeMreB5 was polymerized at room temperature for more than 2 h, which was sufficiently long to obtain a steady state. The baseline was set as the scattering intensity of water. Ionic strength (*IS*) was estimated using the following equation:

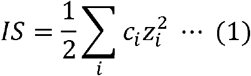

where *c_i_* and *z_i_* are the concentration of the ion species and the charge of the ion, respectively.

## Acknowledgments

We thank Dr. Akihiro Narita (Graduate School of Science, Nagoya University, Japan) and Prof. Katsumi Imada (Graduate School of Science, Osaka University, Japan) for helpful discussions and Mr. Yuhei O Tahara and Ms. Junko Shiomi (Graduate School of Science, Osaka Metropolitan University, Japan) for technical assistances with electron microscopy and SpeMreB expression, respectively.

## Supplemental figures

A PDF file containing Fig. S1-5 is prepared as the file set for supplemental figures.

