## Supporting figures and legends for "Assembly properties of *Spiroplasma* MreB involved in swimming motility"

* To whom correspondence should be addressed:

**Figure S1. Negative-staining EM observation of SpeMreB3 and SpeMreB5 over various pH and ionic strength.** (**A-G**) Negative-staining EM images of 10 μM SpeMreB5 polymerized with 2 mM Mg-ATP at pH (**A-B**) 5, (**C-E**) 7, and (**F-G**) 9 with KCl concentrations of (**C**) 50, (**A, D,** and **F**) 200, and (**B, E,** and **G**) 400 mM. (**H-L**) Negative-staining EM images of 10 μM SpeMreB3 polymerized with 2 mM Mg-ATP at pH (**K**) 5, (**H-J**) 7, and (**L**) 9 with KCl concentrations of (**H** and **K-L**) 50, (**I**) 200, and (**J**) 400 mM. (**M-R**) Negative-staining EM images of (**M-N**) 10 and (**R**) 5 μM SpeMreB5 and (**O-Q**) 10 μM SpeMreB3 incubated in the absence of Mg-ATP at pH (**M** and **O**) 5, (**P** and **R**) 7, and (**N** and **Q**) 9 with (**M-Q**) 50 and (**R**) 200 mM KCl. (**S**) Negative-staining EM image of 10 μM SpeMreB5 polymerized in the absence of Mg-ATP in previously defined standard buffer (composed of 20 mM Tris-HCl pH 7.5 and 100 mM KCl) (1). (**T**) Negative-staining EM image of 10 μM SpeMreB5 ΔC9 polymerized in the absence of Mg-ATP at pH 7 with 10 mM KCl. Scale bars are indicated in each panel.

**Figure S2. Surface potential maps of MreB family proteins.** The coulombic electrostatic potential is indicated by a color gradient from blue (10 kcal/mol/e) to red (-10 kcal/mol/e); namely blue, white, and red regions indicate positively charged, uncharged, and negatively charged regions, respectively. The structural models of double-stranded filament were created by fitting four MreB molecules to each subunit of a double-stranded filament structure of CcMreB (PDB: 4CZJ). (**A-C**) Surface potential maps on the membrane side of double-stranded filament model of

SciMreB5 AMPPNP (PDB: 7BVY) at pH (**A**) 5, (**B**) 7, and (**C**) 9. The positions of facing subdomains (IA and IB) are labeled for the left side subunits. A circle with dashed line indicates the region where is possibly occupied by the positively charged C-terminus, which is not visualized in the reference structure. (**D-F**) Surface potential maps of protofilaments and double-stranded filament models of (**D**) SpeMreB3 (PDB: 7E1G), (**E**) TmMreB (PDB: 1JCG), and (**F**) EcMreB (the monomer was modeled by AlphaFold2 (2)) at pH 7. The protofilament model of EcMreB (front and back side views) were created by fitting two EcMreB molecules to each subunit of a protofilament of a CcMreB double-stranded filament (PDB: 4CZJ). A circle with dashed line on the membrane side of SpeMreB3 double-stranded filament model indicates the region that is possibly occupied by the amphipathic N-terminal helix, which is not visualized in the reference structure. Lysine residues of SpeMreB3 are di-methylated (1), weakening the positive charge of lysine residues. Therefore, the di-methylated lysine residues were replaced with the most probable rotamers of lysine in Dunbrank's rotamer library (3).

**Figure S3. Paracrystal formation and disaggregation dynamics of SpeMreB5.** For time-course light scattering, representative traces from three repeated assays for each condition are shown. (**A-B**) Negative-staining EM images of 10 μM SpeMreB5 polymerized with 2 mM (**A**) Mg-AMPPNP and (**B**) Mg-ADP at pH 7 with 50 mM KCl. Scale bars are indicated in each panel. (**C**) Assembly dynamics of 10 μM SpeMreB5 in the presence of (green) 2 mM Mg-ATP, (light green) Mg-AMPPNP, and (orange) Mg-ADP at pH 7 with 10 mM KCl. (**D**) Dynamics of SpeMreB5 paracrystal formed in the absence of nucleotides. SpeMreB5 with 10 μM concentration was assembled in the absence of nucleotides at pH 7 with 10 mM KCl (dotted line on the left graph). After several minutes, 2 mM Mg-ATP was added, and time-course light scattering was measured (solid line in the right graph). (**E**) Assembly dynamics of 10 μM SpeMreB5 polymerized with 2 mM Mg-ATP at pH 7 with (green) 10 and (light green) 20 mM KCl measured by light scattering. (**F**) Negative-staining EM of 10 μM SpeMreB5 polymerized for 2 min at pH 7 with 20 mM KCl in the presence of 2 mM Mg-ATP. The image was taken at another field of the same EM grid as that for Fig. 3D. Scale bar is indicated in the panel. (**G**) Unnormalized plots of Fig. 4A which shows steady-state light scattering of 10 μM SpeMreB5 polymerized with 2 mM Mg-ATP over the range of KCl concentration at pH (ocher) 5 and (green) 7. Bars indicate S.D. from three independent measurements.

**Figure S4. Sedimentation assays of SpeMreB5.** Each sample was incubated in buffer S, centrifuged, and analyzed using SDS-PAGE, as previously described (1). All reactions were conducted in the presence of 2 mM ATP. For divalent cation-free conditions, 1 mM EDTA-NaOH pH 8.0 was added to avoid effects from contaminating the amounts of multivalent cations. (**A-D**) Sedimentation assays of SpeMreB5 (**A** and **C-D**) WT and (**B**) ΔC9 polymerized with 2 mM (**A-B**) Mg-ATP, (**C**) Ca-ATP, and (**D**) ATP (divalent cations free) over the range of SpeMreB5 concentration. (**E**) Quantified precipitation amounts of sedimented SpeMreB5. Precipitated SpeMreB5 was resuspended in water equivalent amount to the sample, and the resulting concentrations were plotted over the total SpeMreB5 concentrations with linear fitting. Error bars indicate S.D. from three repeated measurements. Critical concentrations were estimated as the x-intercept of each linear fit and summarized in Table 1. (**F-G**) Sedimentation assay of 3 μM SpeMreB5 over the range of (**F**) $MgCl_2$ and (**G**) $CaCl_2$ concentrations. (**H**) Quantified precipitation amounts of sedimented SpeMreB5. Precipitated SpeMreB5 was resuspended in water equivalent amount to the sample, and the resulting concentrations were plotted over the divalent cation concentrations with linear fitting. Error bars indicate S.D. from three repeated measurements.

**Figure S5. Divalent cation dependence of SpeMreB5 paracrystal.** For time-course light scattering, representative traces from three repeated assays for each condition are shown. For divalent cation-free conditions, 1 mM EDTA-NaOH pH 8.0 was added to avoid effects from contaminating amounts of multivalent cations. (**A-B**) Negative-staining EM images of 10 μM SpeMreB5 polymerized in the presence of (**A**) 2 mM Ca-ATP at pH 5 and (**B**) 2 mM ATP at pH 7. KCl concentration was constant for 50 mM KCl. The image of the panel B was taken at another field of the same EM grid as that for Fig. 6B. Scale bars are indicated in each panel. (**C-D**) $Ca^{2+}$-dependent assembly dynamics of 10 μM SpeMreB5 at pH (**C**) 5 and (**D**) 7 with 10 mM KCl measured using light scattering. Polymerization was initiated by adding 2 mM ATP with varying $CaCl_2$ concentrations as indicated in the color scales in the panels. (**E**) Unnormalized light scattering traces of Fig. 6G that shows disassembly dynamics of SpeMreB5 paracrystals induced with divalent cations. (**F**) Time-course light scattering measurements of SpeMreB5 paracrystals disassembly induced by 10 mM HEPES-KOH pH 8.1 (green) and EDTA-NaOH pH 8.0 (purple). The initial paracrystal solutions were prepared by polymerizing 10 μM SpeMreB5 in the presence of 2 mM Mg-ATP in 10 mM HEPES-KOH pH 7.0 with 40 mM KCl. The measurement in which the buffer composition was unchanged are indicated with a dark green line.

### Fig S1

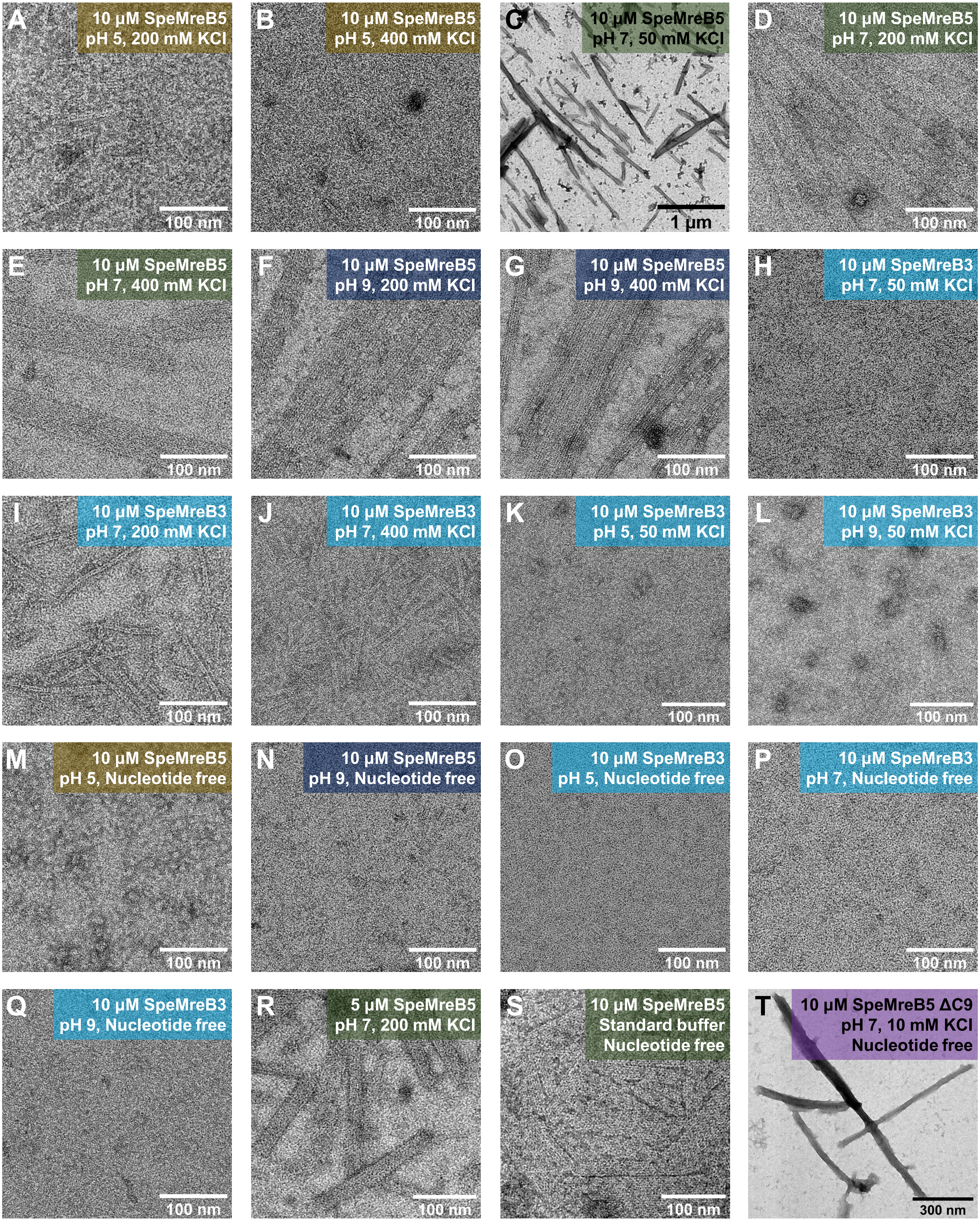
A
10 µM SpeMreB5
pH 5, 200 mM KCl
100 nm
B
10 µM SpeMreB5
pH 5, 400 mM KCl
100 nm
C
10 µM SpeMreB5
pH 7, 50 mM KCl
1 µm
D
10 µM SpeMreB5
pH 7, 200 mM KCl
100 nm
E
10 µM SpeMreB5
pH 7, 400 mM KCl
100 nm
F
10 µM SpeMreB5
pH 9, 200 mM KCl
100 nm
G
10 µM SpeMreB5
pH 9, 400 mM KCl
100 nm
H
10 µM SpeMreB3
pH 7, 50 mM KCl
100 nm
I
10 µM SpeMreB3
pH 7, 200 mM KCl
100 nm
J
10 µM SpeMreB3
pH 7, 400 mM KCl
100 nm
K
10 µM SpeMreB3
pH 5, 50 mM KCl
100 nm
L
10 µM SpeMreB3
pH 9, 50 mM KCl
100 nm
M
10 µM SpeMreB5
pH 5, Nucleotide free
100 nm
N
10 µM SpeMreB5
pH 9, Nucleotide free
100 nm
O
10 µM SpeMreB3
pH 5, Nucleotide free
100 nm
P
10 µM SpeMreB3
pH 7, Nucleotide free
100 nm
Q
10 µM SpeMreB3
pH 9, Nucleotide free
100 nm
R
5 µM SpeMreB5
pH 7, 200 mM KCl
100 nm
S
10 µM SpeMreB5
Standard buffer
Nucleotide free
100 nm
T
10 µM SpeMreB5 ΔC9
pH 7, 10 mM KCl
Nucleotide free
300 nm

### Fig S2

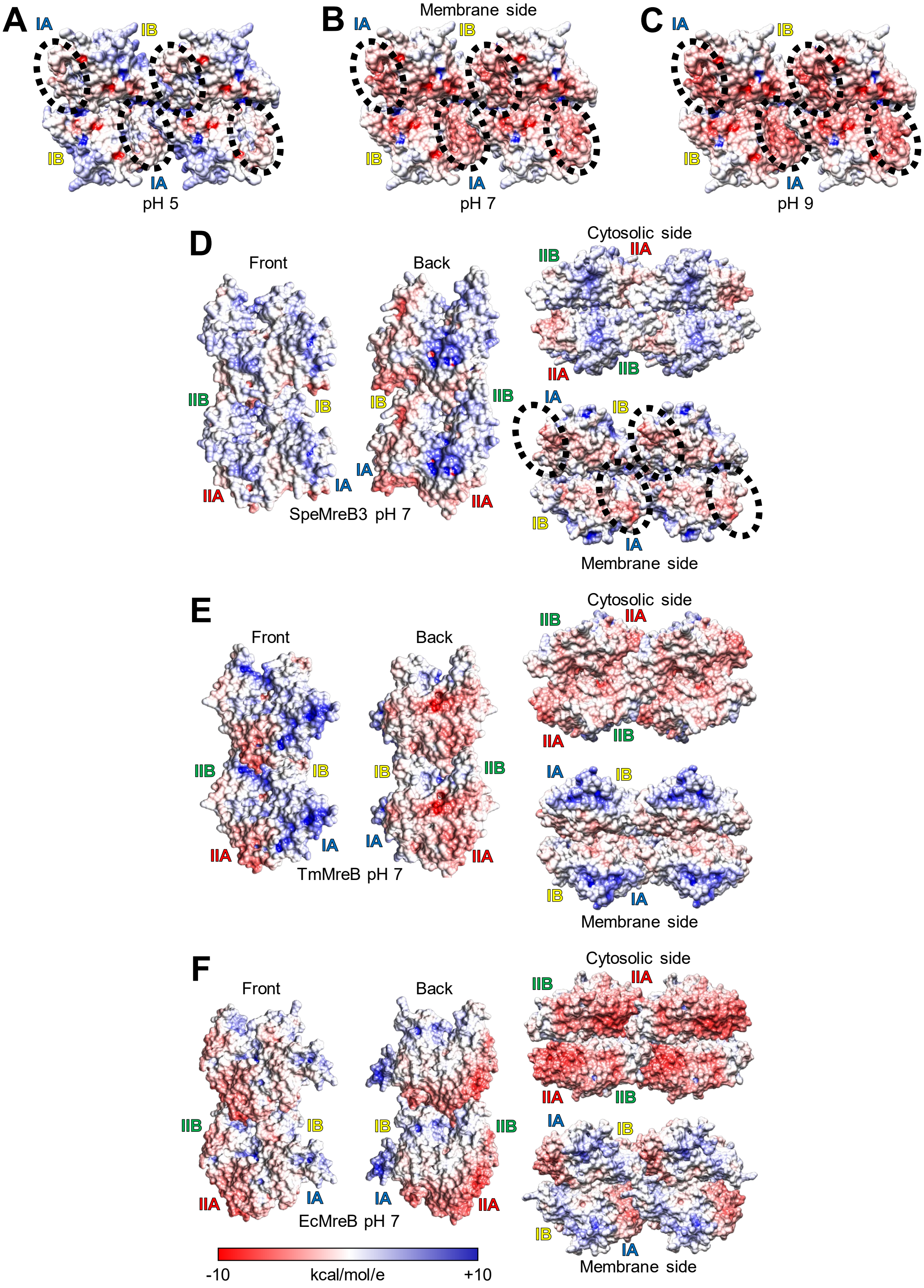
A
IA
IB
IB
IA
pH 5
B
Membrane side
IA
IB
IB
IA
pH 7
C
IA
IB
IB
IA
pH 9
D
Front
Back
Cytosolic side
IIB
IIA
IIA
IIB
IIB
IB
IB
IIB
IA
IB
IA
IA
IIA
IIA
IB
IA
SpeMreB3 pH 7
Membrane side
E
Front
Back
Cytosolic side
IIB
IIA
IIA
IIB
IIB
IB
IB
IIB
IA
IB
IA
IA
IIA
IIA
IB
IA
TmMreB pH 7
Membrane side
F
Front
Back
Cytosolic side
IIB
IIA
IIA
IIB
IIB
IB
IB
IIB
IA
IB
IIA
IA
IA
IIA
IB
IA
EcMreB pH 7
Membrane side
-10
kcal/mol/e
+10

### Fig S3

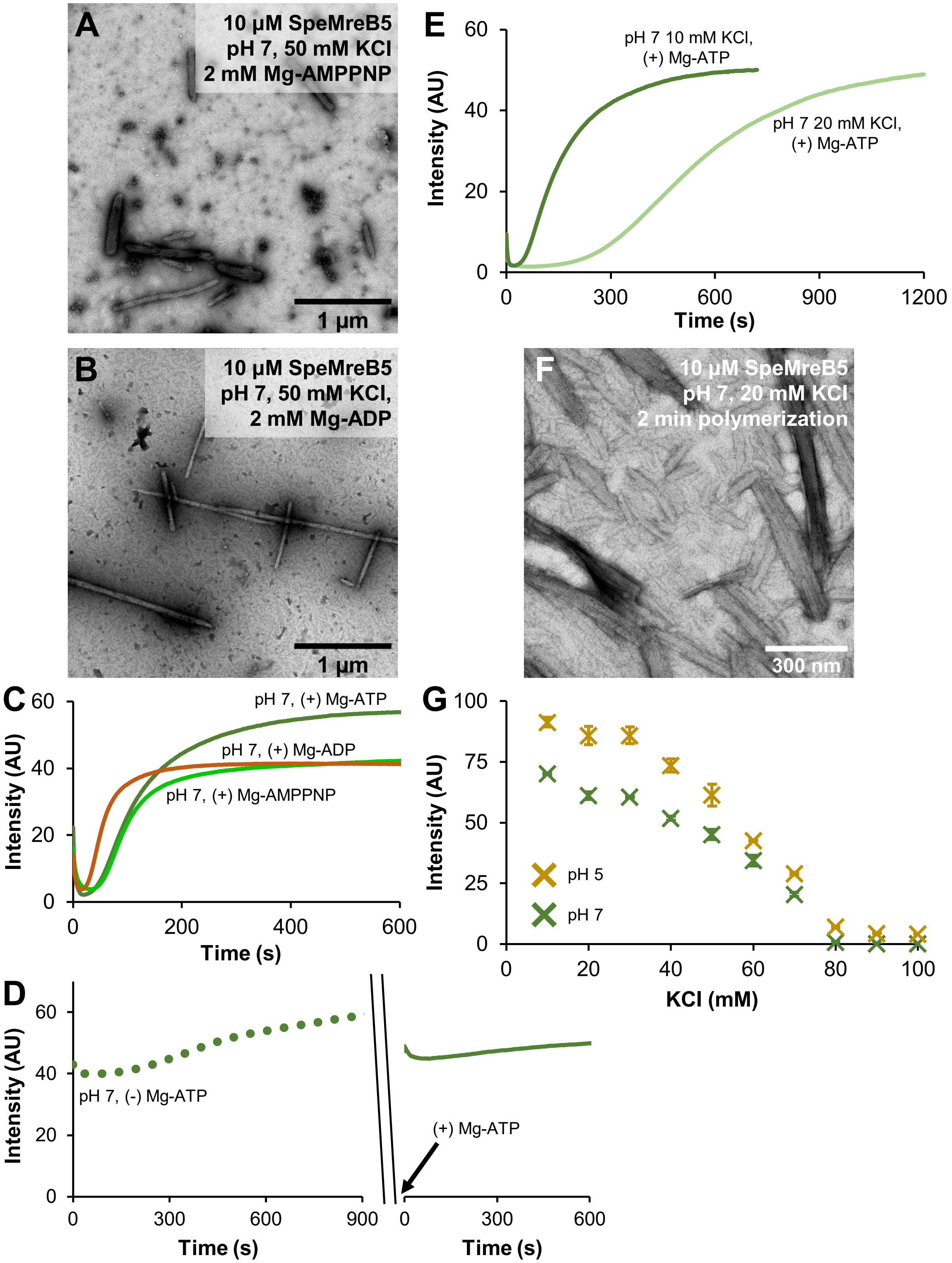
A
10 µM SpeMreB5
pH 7, 50 mM KCl
2 mM Mg-AMPPNP
1 µm
B
10 µM SpeMreB5
pH 7, 50 mM KCl,
2 mM Mg-ADP
1 µm
C
pH 7, (+) Mg-ATP
pH 7, (+) Mg-ADP
pH 7, (+) Mg-AMPPNP
Intensity (AU)
Time (s)
D
pH 7, (-) Mg-ATP
(+) Mg-ATP
Intensity (AU)
Time (s)
Time (s)
E
pH 7 10 mM KCl,
(+) Mg-ATP
pH 7 20 mM KCl,
(+) Mg-ATP
Intensity (AU)
Time (s)
F
10 µM SpeMreB5
pH 7, 20 mM KCl
2 min polymerization
300 nm
G
pH 5
pH 7
Intensity (AU)
KCl (mM)

### Fig S4

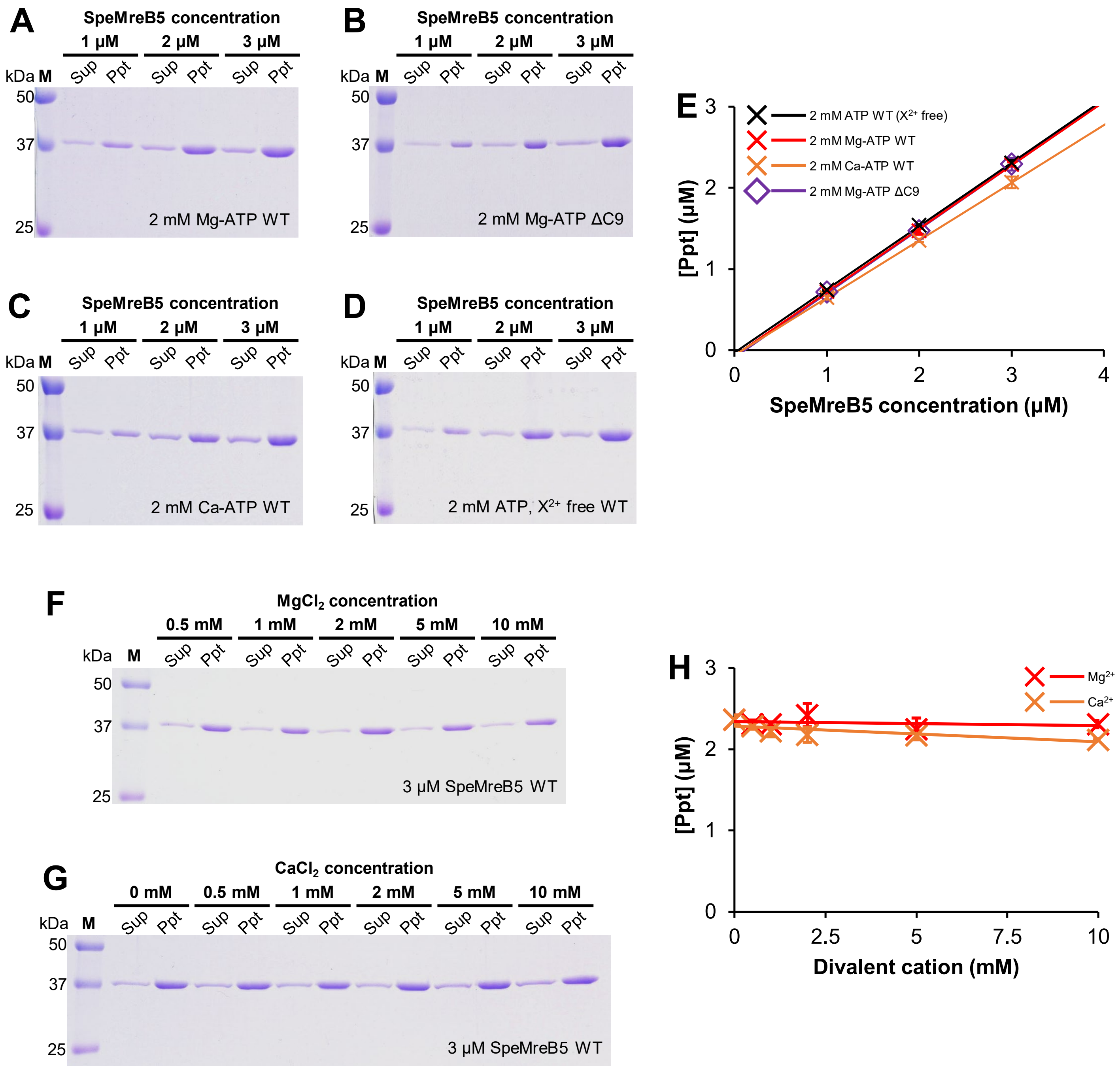
A
SpeMreB5 concentration
1 μM
2 μM
3 μM
kDa M
Sup Ppt Sup Ppt Sup Ppt
50
37
25
2 mM Mg-ATP WT
B
SpeMreB5 concentration
1 μM
2 μM
3 μM
kDa M
Sup Ppt Sup Ppt Sup Ppt
50
37
25
2 mM Mg-ATP ΔC9
C
SpeMreB5 concentration
1 μM
2 μM
3 μM
kDa M
Sup Ppt Sup Ppt Sup Ppt
50
37
25
2 mM Ca-ATP WT
D
SpeMreB5 concentration
1 μM
2 μM
3 μM
kDa M
Sup Ppt Sup Ppt Sup Ppt
50
37
25
2 mM ATP, X²⁺ free WT
E
2 mM ATP WT (X²⁺ free)
2 mM Mg-ATP WT
2 mM Ca-ATP WT
2 mM Mg-ATP ΔC9
[Ppt] (μM)
0
1
2
3
4
SpeMreB5 concentration (μM)
F
MgCl₂ concentration
0.5 mM
1 mM
2 mM
5 mM
10 mM
kDa M
Sup Ppt Sup Ppt Sup Ppt Sup Ppt Sup Ppt
50
37
25
3 μM SpeMreB5 WT
G
CaCl₂ concentration
0 mM
0.5 mM
1 mM
2 mM
5 mM
10 mM
kDa M
Sup Ppt Sup Ppt Sup Ppt Sup Ppt Sup Ppt Sup Ppt
50
37
25
3 μM SpeMreB5 WT
H
Mg²⁺
Ca²⁺
[Ppt] (μM)
0
1
2
3
2.5
5
7.5
10
Divalent cation (mM)

### Fig S5

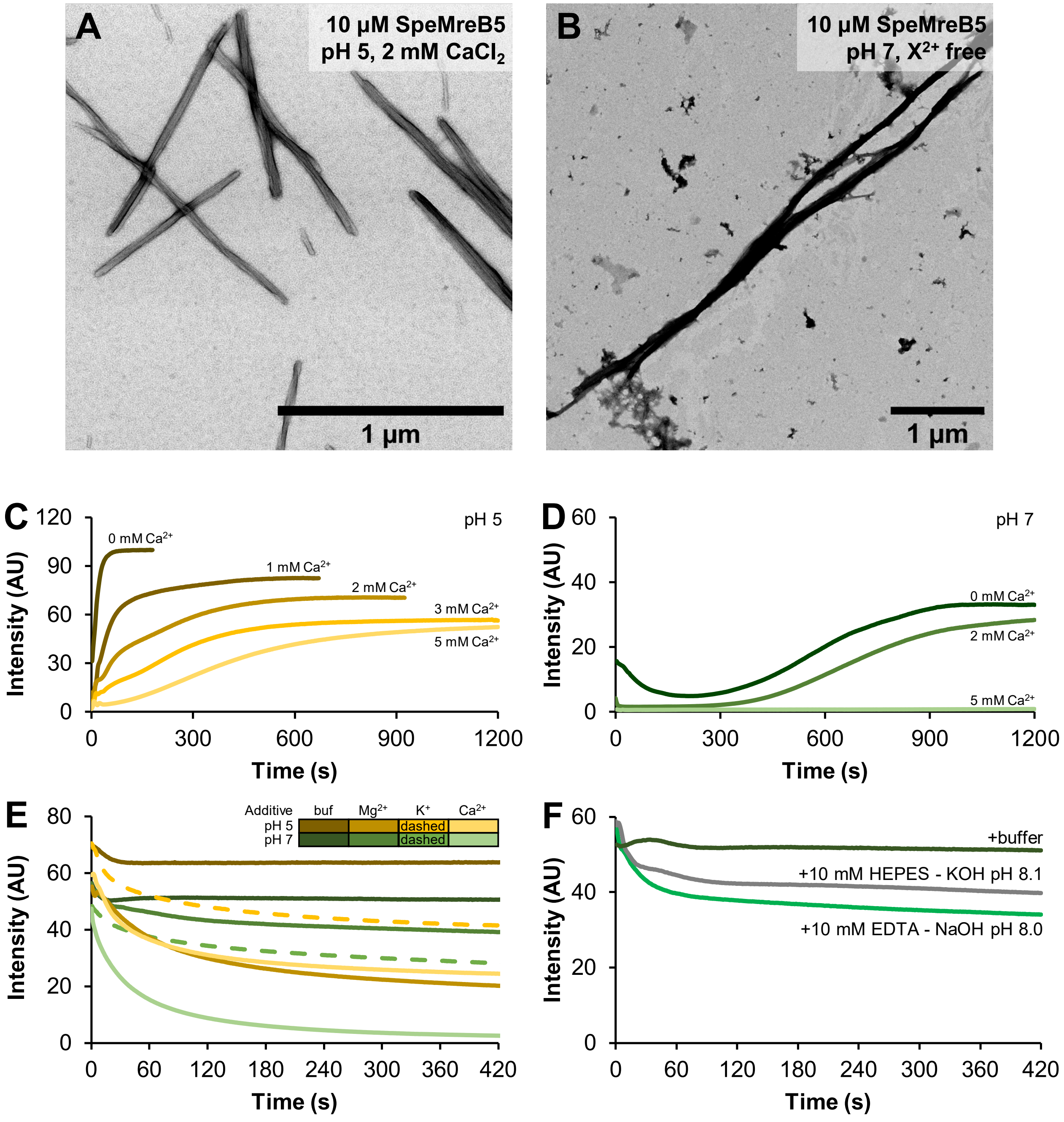

A
10 µM SpeMreB5
pH 5, 2 mM CaCl$_2$
1 µm
B
10 µM SpeMreB5
pH 7, X$^{2+}$ free
1 µm
C
pH 5
0 mM Ca$^{2+}$
1 mM Ca$^{2+}$
2 mM Ca$^{2+}$
3 mM Ca$^{2+}$
5 mM Ca$^{2+}$
Intensity (AU)
Time (s)
D
pH 7
0 mM Ca$^{2+}$
2 mM Ca$^{2+}$
5 mM Ca$^{2+}$
Intensity (AU)
Time (s)
E
Additive buf Mg$^{2+}$ K$^{+}$ Ca$^{2+}$
pH 5 dashed
pH 7 dashed
Intensity (AU)
Time (s)
F
+buffer
+10 mM HEPES - KOH pH 8.1
+10 mM EDTA - NaOH pH 8.0
Intensity (AU)
Time (s)
